# Biophysical properties of IgD determine thresholds for self-tolerance and selection into germinal centers

**DOI:** 10.64898/2026.07.02.735902

**Authors:** Lachlan P. Deimel, Ryan A. Brady, Andriy Goychuk, Gabriela S Silva Santos, Clara Uhe, Andrew J. MacLean, Brianna Hernandez, Shuai Zong, Laurine Binet, Mary Tenuta, Dennis Shaefer-Babajew, Zeliha Kilic, Anna Gazumyan, Thiago Y. Oliveira, Harald Hartweger, Arup K. Chakraborty, Scott C. Blanchard, Michel C. Nussenzweig

## Abstract

Immunoglobulin D (IgD) is among the most conserved antibody isotypes, found in virtually all jawed vertebrates^1^. Unlike other isotypes, IgD contains an unusually long hinge region of up to 160 amino acids that connects the constant and variable regions. Its expression pattern is also conserved; IgD is co-expressed with IgM on transitional and mature naïve B cells. However, the function of IgD has remained enigmatic since its discovery in 1965^2,3^. Here we present and test a biophysical model positing that IgD increases the entropic cost of bivalent antigen binding. Single-molecule measurements revealed that the antigen-binding arms of IgD are substantially more dynamic than those of IgM, suggesting that cell surface IgD would be energetically penalized in bivalent antigen binding. Consistent with the model and biophysical data, we find that the long hinge compromises antigen capture by IgD B cell receptors (BCRs) compared to IgM BCRs. To determine how the difference in antigen binding impacts immunity, we produced mice that express only IgM and IgD, exclusively IgM or IgD, or IgD with a truncated hinge region. The data indicate that the increased entropic cost of antigen binding imposed by the IgD hinge attenuates negative selection by self-antigen while increasing the affinity-based threshold for positive selection into the germinal center (GC). Together the results indicate that IgD functions physiologically to desensitize B cells to antigen, thereby expanding the B cell repertoire while optimizing affinity-based selection into the GC.

---

B cells must interpret and respond to an extraordinarily complex antigenic environment. Antigen is sensed by a cell’s unique BCR, which is composed of any one of several different membrane-bound isotypes of immunoglobulin (Ig) associated with Igɑ-Igβ signal transducers^4,5^. Encounters between BCR and cognate antigen can initiate a series of different responses including cell death, tolerance, survival or activation, and these decisions depend on context, developmental stage and interaction with other immune cells^6,7^. For example, self-antigen recognition by developing B cells results in elimination of autoreactive clones by deletion, anergy or receptor editing^8–13^. In contrast, mature naïve B cells can respond to antigen by undergoing clonal expansion and producing antibodies that are essential for pathogen clearance^14^. Precisely how the different isotypes of the BCR help instruct these distinct responses is poorly understood.

Developing B cells in the bone marrow express only a single Ig isotype, IgM. Upon leaving the bone marrow, transitional and mature naïve B cells use alternative splicing to co-express IgM and IgD BCRs at an average ratio of 1:20^15,16^. The two are unique among Ig isotypes in their capacity for co-expression and in the mechanisms that regulate their dynamic cell surface levels^16,17^. In addition, they share the same signal transducers, Igɑ-Igβ^4^. Although, IgD knockout mice and heterozygous IgD-deficient humans show no gross defects in B cell development^18–21^, increased IgD expression is associated with B cell autoreactivity and anergy^12,22,23^.

One of the key differences between IgM and IgD is the hinge region separating the Ig constant from the variable region (Fig. 1a). IgM lacks a classical hinge domain, and structural evaluation by electron microscopy suggests that its fragment antigen binding (Fab) arms are relatively constrained^24,25^. In contrast, IgD has an exceedingly long linker region of 35 and 64 residues in mice and humans^1^, respectively. AlphaFold-Multimer modelling indicates that the IgD linker is highly disordered (Supplementary data file 1)^26^. In contrast, synchrotron X-ray scattering and molecular dynamics modeling suggest that IgD exists primarily in a T shaped structure^27^, and crystal structures of the CH1 domain reveal a conformationally restricted upper hinge that could contribute to stabilizing a T shaped structure^28^. However, the precise molecular dynamic properties of IgD, and how they might impact B cell physiology, remain to be determined.

**Fig. 1:**
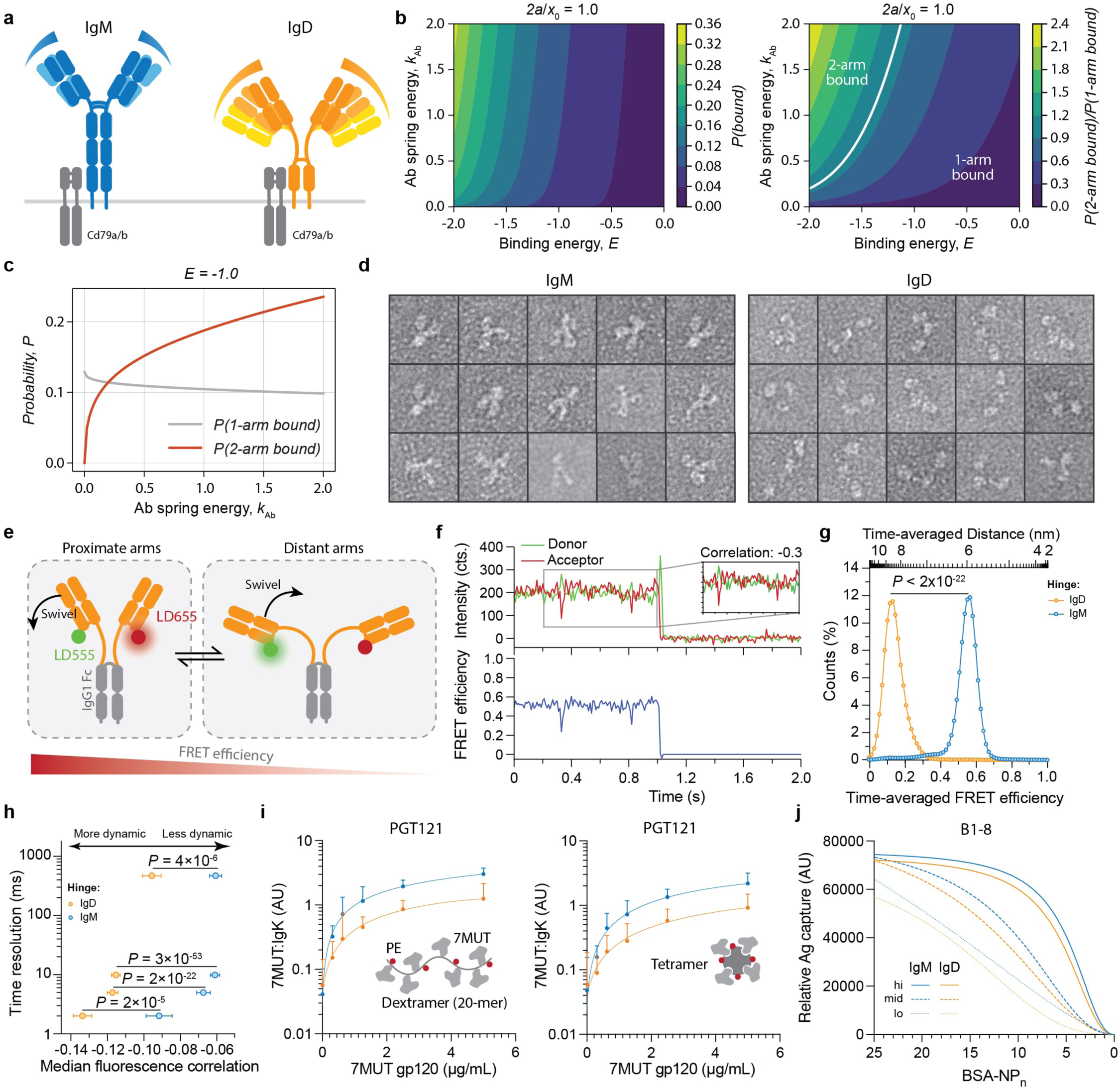
Biophysical properties of IgM and IgD, and their implications for antigen binding. **a,** Diagrammatic representation of the mouse IgM and IgD BCR. **b,c,** Plots showing the calculated probability of antibody binding as a function of Fab arm affinity/binding energy and spring energy/hinge rigidity. Calculations are described in Supplementary Document 1. **d,** Representative particle images of purified mouse IgM and IgD obtained by negative stain electron microscopy (50 000 x magnification). **e,** Graphical depiction of single molecule FRET strategy, labeling the C-terminal A4 tag with CoA–LD555 and CoA–LD655 with AcpS enzyme. **f,** Example single-molecule fluorescence (top) and FRET (bottom) trace collected at 10 ms time-resolution. Traces show rare, transient, anti-correlated excursions (inset) indicating fast dynamics. cts.; photon counts. **g,** Population histograms showing the time-averaged FRET efficiencies and inter-dye distance for data collected at 10 ms time resolution for 4409 IgG1:IgD molecules and 4372 IgG1:IgM molecules. *P*-value determined by Wilcoxon rank sum test. **h,** Correlation coefficients of single-molecule donor and acceptor fluorescence at varying exposure times (500 ms, 10 ms, 5 ms and 2 ms) were used to obtain the median for each measurement. Data points and error bars show the mean and standard error of median correlation coefficients, respectively, as determined from 1000 bootstrap resamples of a combined dataset from at least 5 independent measurements. *P*-values are indicated, as determined by two-sided Wilcoxon rank sum test. **i,j,** HEK293T cells were transiently transfected with plasmids encoding a complete mouse BCR: plasmids encoded the light chain, Cd79a/b, and either IgM or IgD heavy chain. 16 h after transfection, antigen binding was assessed using fluorescently-labelled antigen. **i,** Cells were transfected with anti-gp120 (7MUT) antibody clone, PGT121, expressed as either IgM or IgD BCR. Cell capture of dextramerized (20-mer) or tetramerized 7MUT antigen was recorded and normalized to the BCR expression. Dots indicate the mean Ag-IgK ratio, with bars denoting standard deviation. Data were acquired from at least 150 cells per group. **j,** NP-binding clone, B1-8, variants of different affinities (B1-8^hi^: k_D_ = 200, B1-8^mid^: k_D_ = 2000 nM and B1-8l°: k_D_ = 8000 nM)^74^ was displayed by cells. Capture of differentially nitrophenylated albumin was titrated.

To understand how Fab arm dynamics might impact antigen binding, we developed a theoretical model that considers the biophysical relationship between Fab–antigen affinity and hinge flexibility. The latter is modelled as a spring, with spring stiffness (or energy associated with deforming it) being a proxy of conformational rigidity (Fig. 1b,c; Supplementary document 1). The model accounts for antigen topology, defined here as the spatial arrangement of the epitopes—specifically, the distance between two identical epitopes (2a) relative to the preferred separation between the two Fab arms of the BCR (x_0_). Since both BCR and antigen are somewhat diffuse on a liquid membrane, and able to sample all possible topologies, we assume 2a/x_0_ = 1.0^29–31^. Our biophysical calculations predict that the probability of a BCR being in a two arm-bound state increases with hinge stiffness irrespective of antibody affinity (Fig. 1b,c).

To directly visualize the Fab arms of IgM and IgD, we examined them by negative-stain electron microscopy, which revealed the expected variation in IgD Fab arm positioning compared with IgM (Fig. 1d)^27,32^.

To assess the dynamic motion of the Fab arms in real-time, we performed single-molecule Förster resonance energy transfer (smFRET) analyses using a prism-based total internal reflection fluorescence (pTIRF) imaging platform. We engineered chimeric Ig wherein the hinge and Fab arms of murine IgD or IgM were transplanted onto the murine IgG1 Fc (Fig. 1e). The light chains were *C*-terminally tagged with the A4 peptide, which was directly labelled with coenzyme A (CoA)-conjugated dyes using holo-[acyl-carrier-protein] synthase (AcpS; refer to Methods for labelling details)^33^. We used FRET donor, LumiDyne (LD)555-CoA, and acceptor, LD655-CoA. The *C*-termini of the heavy chains were directly biotinylated using biotin ligase (BirA) through introduction of an Avi-tag to enable surface immobilization in a biotin-polyethylene glycol (PEG) passivated quartz imaging chamber through a streptavidin bridge. Fluorescence (Fig. 1f, top panel) and FRET traces (Fig. 1f, bottom panel) show relatively stable behaviour, with short-lived anti-correlated excursions occasionally observed at high imaging rates (2–10 ms) (Fig. 1f, inset). This indicates rapid Fab arm dynamics (>500 s^-1^). Population FRET efficiency distributions of the two constructs revealed that the intramolecular time-averaged distance between the *C*-termini of the Fabs, 〈*r*〉, is significantly larger for IgD (FRET = 0.11; 〈*r*〉 = 8.8 nm) than for IgM (FRET = 0.55; 〈*r*〉 = 6.0 nm) (*P* < 2 x 10^-22^; Fig. 1g). To compare Fab arm dynamics between constructs, we calculated the anticorrelation of donor and acceptor fluorescence across a range of time-resolutions (Fig. 1g; Supplementary data fig. 1). Here, the greater the anticorrelation, the more motile the Fab arms^34^. We observed significantly more negative median correlation for IgD than for IgM across all time-resolutions tested (2 ms; *P* = 2 x 10^-5^; 5ms, *P* = 2 x 10^-22^; 10 ms, *P* = 3 x 10^-53^; 500 ms, *P* = 4 x 10^-6^). These data demonstrate that the Fab arms of IgD are both more extended and significantly more dynamic than those of IgM.

The biophysical and modelling data predict that cell surface IgM captures antigen more efficiently than IgD, due to differences in the entropic cost of bivalent binding. To test this idea, we developed an *in vitro* system using HEK293T cells to display mouse IgM or IgD along with Igɑ/Igβ (Supplementary data fig. 2a,b). Anti-HIV-1 antibodies with 3 different affinities (k_D_: 1.2nM, 12 µM and > 36 µM) were expressed. A fluorescently labelled HIV-1 gp120 antigen (7MUT)^35^ was titrated as a tetramer or dextramer (20-mer), and binding was normalized to surface BCR density (Fig. 1i; Supplementary data fig. 2c,d). The IgM BCR captured more antigen compared to IgD across a diverse range of BCR affinities, antigen concentrations and degrees of multimerization (Fig. 1i; Supplementary data fig. 2c,d). A similar experiment was performed using 4-hydroxy-3-nitrophenylacetyl (NP)-specific antibody variants, B1-8, which showed a similar isotype-dependent effect on antigen capture (Fig. 1j).

To confirm that differential antigen capture was a function of the length of the hinge, we deleted the amino acids corresponding to the linker exon of IgD to produce a hinge region that resembles that of IgM (IgD^Trunc^; Supplementary data fig. 3a). When expressed as an anti-NP BCR, B1-8 IgD^Trunc^, antigen capture was equivalent to IgM (*P* = 0.46) and superior to IgD (*P* < 0.0001; Supplementary data fig. 3b). Thus, when expressed on the surface of transfected cells, the hinge of cell surface IgD renders it inferior to IgM for antigen capture.

Our model also predicts that some antigen display topologies disfavour stiffer BCRs, that is, where 2a >> x_0_ (Supplementary data fig. 4a). For example, if two copies of the antigen are in fixed distant positions, IgD would be at an advantage because it has greater and more variable reach. We tested this idea by randomly immobilizing NP-conjugated bovine serum albumin (BSA) to a solid surface and measured the binding of B1-8^hi^ expressed as soluble monomeric IgM or IgD by enzyme-linked immunosorbent assay (ELISA). Consistent with the biophysical calculations, soluble B1-8 expressed as IgD bound randomly immobilized NP more effectively than IgM (Supplementary data fig. 4b)^32^. This feature may be relevant for particulates such as viruses that display repetitive antigens spaced at fixed distances that could prevent bivalent binding by IgM but not IgD.

To test the hypothesis that cell surface expression of IgD would be less sensitive to self- or foreign antigens compared to IgM, we produced mice that express IgM and IgD (MD-only), exclusively IgM (M-only^36^), or exclusively IgD (D-only; Supplementary data fig. 5a, Fig. 2a). B cells from MD-only mice express both isotypes, whereas their M- or D-only counterparts are restricted to IgM or IgD expression, respectively. Hence, M-only and D-only mice deviate from native surface BCR regulation in the immature (BCR^+^CD23^-^), transitional (CD93^+^) and naïve (CD43^-^CD93^-^CD23^+^) B cell compartments where these isotypes are co-expressed (Fig. 2b; Supplementary data fig. 5b–d). In addition, B cells in MD-, M- or D-only mice cannot undergo class-switch recombination since C_γ3_–C_ɑ_ was removed.

**Fig. 2:**
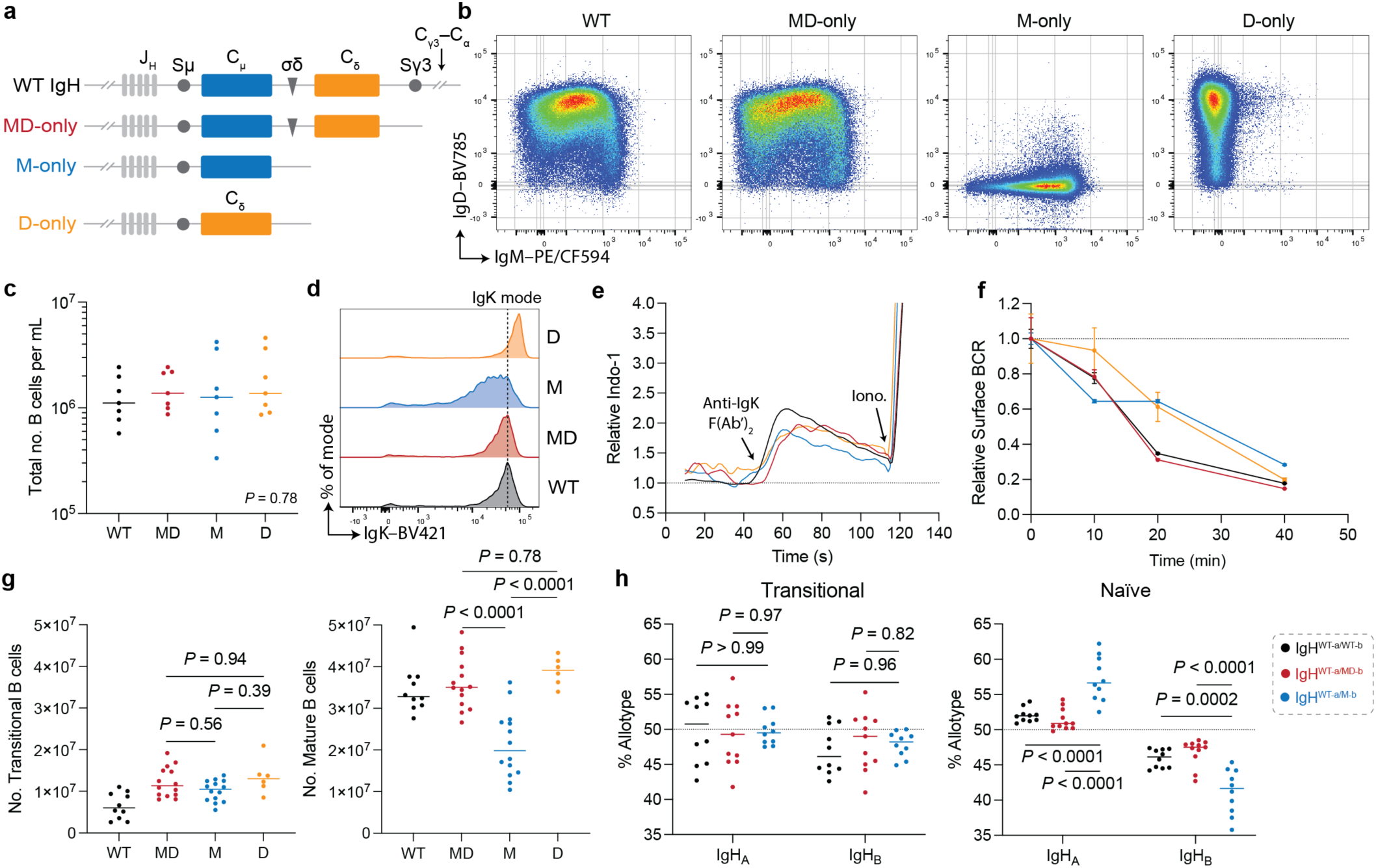
Expression of IgD during peripheral development is permissive of B cell maturation. **a,** Diagrammatic representation of the *Igh* loci from WT, MD-, M- and D-only models. S = class recombination switch region, C = constant region gene fragment, and σδ = alternative splice site. **b,** Flow cytometry plots show the IgM and IgD expression on splenic mature B cells in the transgenic mouse models. **c,** Plot shows the total number of B cells in the blood. **d,** Histogram of surface IgK expression on mature naïve B cells. **e,** Traces show cellular calcium flux over time by primary B cells in response to anti-mouse IgK F(ab’)_2_ followed by ionomycin. **f,** Graph shows surface BCR, which was monitored longitudinally by flow cytometry after culture, triggering internalization with anti-mouse IgK F(ab’)_2_ (*n* = 4 per group). **g,** Absolute number of splenic transitional and mature naïve B cells were quantified by flow cytometry in 5-week-old mice. **h,** WT, MD- and M-only mice (which bear the *Ighb* allotype) were crossed with discordant *Igha* mice. Plot shows allotype expression by transitional and mature naïve B cells. **c,g,h,** Data are pooled from at least two independent experiments. Dots represent data from a single animal (*n* = 6–14 per group), and bars show the median. Data were compared via ANOVA and post-hoc Tukey’s multiple comparison test. *P*-values are indicated.

Flow cytometry showed normal B cell development in the bone marrow (Fig. 2c, and Supplementary data fig. 5e–g). As expected, the level of cell surface BCR expression in MD-only mice was similar to the wild-type (WT), which was higher than M-only, and lower than in D-only B cells (Fig. 2d)^22^. To confirm that B cells retained their transcriptional program despite the modifications to the *Igh* loci, we performed single-cell transcriptomic analysis on developing and mature B cells (Supplementary data fig. 6). We found no gross differences in the transcriptional programs of mature naïve B cells between strains (Supplementary data fig. 6). Finally, IgVH and IgVL sequencing of the naïve B cell repertoires showed no gross variance in the V-gene usage, proportion of IgK/IgL sequences, average heavy-chain complementary determining region (CDRH)3 length or in the frequency of charged, hydrophobic or aromatic residues (Supplementary data fig. 7).

To examine BCR function, we measured calcium flux in response to receptor crosslinking (Fig. 2e). Primary splenic B cells were stimulated with anti-IgK F(ab’)_2_ and intracellular calcium was measured over time. Consistent with the work of others, all 3 types of mutant B cells were like WT with respect to induction time and magnitude of flux (Fig. 2e)^37,38^. BCR internalization as measured by flow cytometry after anti-IgK F(ab’)_2_ crosslinking *in vitro* was also found to be equivalent between the mutant B cells and the WT control (Fig. 2f).

Under physiologic circumstances, IgD is first expressed when B cells emerge from the bone marrow as transitional cells that undergo negative selection against peripheral self-antigens^39–41^. IgD expression persists in mature naïve B cells that participate in immune responses to foreign antigens. Our theoretical model and biophysical measurements predict that expression of IgD would render B cells less prone to peripheral negative selection in the transition between Immature and mature B cell compartments compared with IgM^42^. Consistent with the predictions, M-only mice accumulated 56% fewer mature naïve B cells compared with MD-only mice (*P* < 0.0001; Fig. 2f). Conversely, D-only mice exhibited normal numbers of mature naïve B cells (Fig. 2g; Supplementary data fig. 8a).

To confirm the relative disadvantage of IgM expression for B cell accumulation in the mature naïve compartment, we crossed WT, MD- and M-only mice (all of which possess the *Ighb* locus) with allotypically discordant WT *Igha* mice. Heterozygous mice maintain allelic exclusion and their B cells express either WT-IgH_A_ or Mutant-IgH_B_ (Supplementary data fig. 8b). Whereas mature naïve B cells expressing IgH_A_ or IgH_B_ alleles were evenly distributed in WT:WT and WT:MD mice, the WT IgH_A_ allele dominated the mature naïve compartment in WT:M mice (Fig. 2h; Supplementary data fig. 8c). To exclude the possibility that the difference in allotype representation was due to altered gene expression by the removal of C_γ3_–C_ɑ_ from the *Igh* locus, we produced mixed MD-only:M-only bone marrow chimeras (Supplementary data fig. 8c). Notably, there were significantly fewer M-only than MD-only B cells in the mature naïve compartment in the chimeras, again indicating that IgD expression was permissive of naïve B cell accumulation (*P* < 0.0001; Supplementary data fig. 8d,e). We hypothesize that expression of IgD reduces sensitivity to self-antigen, allowing B cells to proceed through this self-reactivity checkpoint and accumulate in the mature naïve compartment^42^.

To determine whether IgM and IgD expression alters sensitivity to autoantigens, we examined mice that express two different *Igk* alleles: a pre-arranged self-reactive anti-H2^b^ 3-83 V_k_–J_k_ on one allele^43^, and human C_k_ (huC_k_)^44^ on the other allele. Previous work showed that small pre-BII cells default to expressing the pre-arranged 3-83 IgK but if the Ig heavy chain pairing is non-productive or autoreactive they undergo secondary V(D)J rearrangement that deletes 3-83 IgK resulting in a disproportionate number of B cells that express huC_k_ or the Ig lambda light chain (IgL; Fig. 3a)^44^. The remaining self- or polyreactive 3-83 IgK expressing cells are trapped in the transitional compartment as reflected in an increased proportion of huC_k_ expressing cells and a corresponding decrease of these cells in the mature naïve compartment (Fig. 3b–d).

**Fig. 3:**
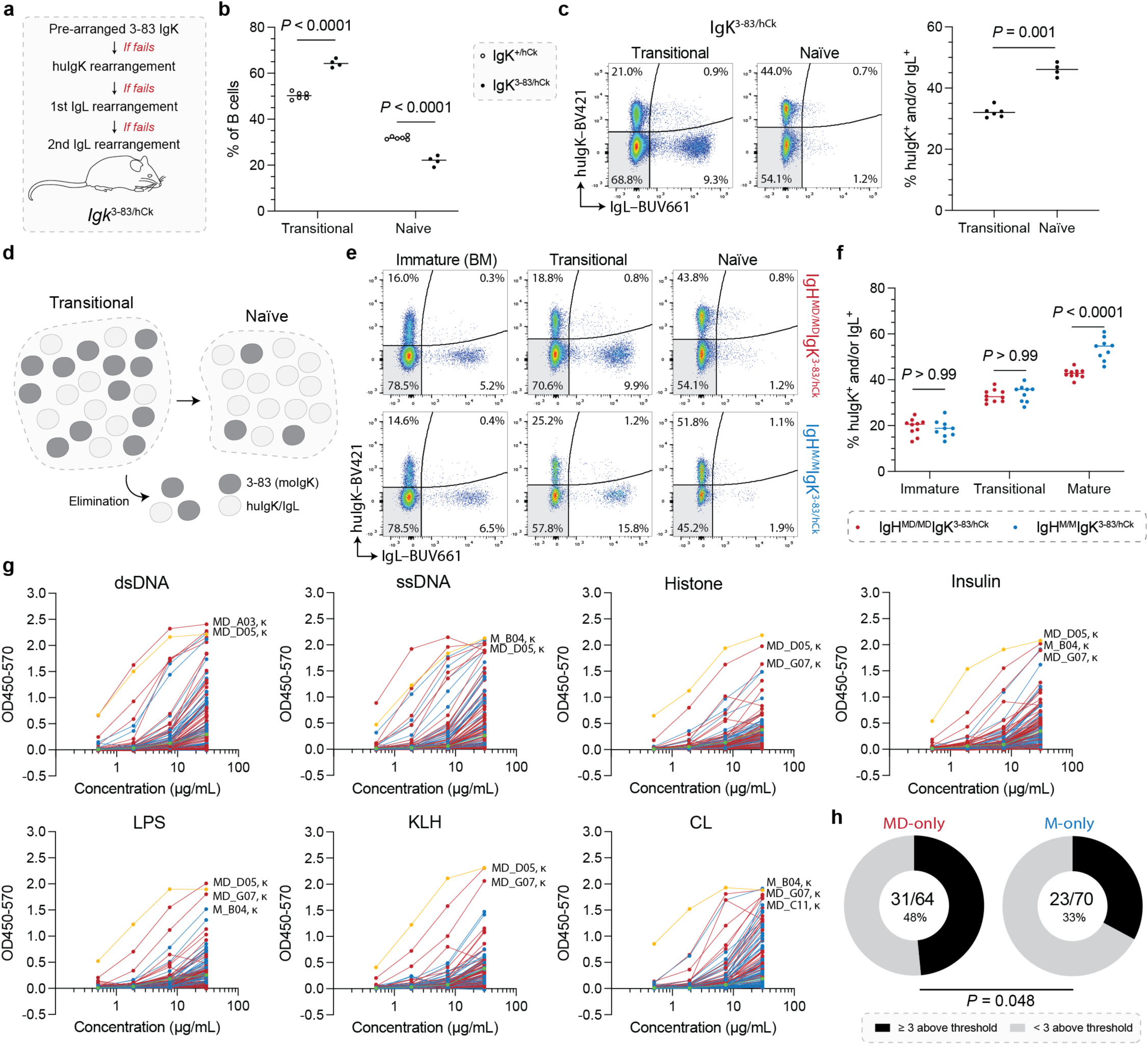
IgD expression tolerizes the naïve B cell repertoire. **a,** Depiction of the light chain rearrangement hierarchy in developing B cells bearing *Igk*^3-83/hCk^ alleles. **b,c,** Mice with a wild-type *Igh* loci were examined. **b,** Plots show the proportion of transitional and naïve B cells in the spleens of 5-week-old *IgK*^+/hCk^ (open circles) and *IgK*^3-83/hCk^ (closed) mice. **c,** Representative flow cytometry plots and dot plot showing the expression of huIgK and IgL in the transitional and naïve compartments of *IgK*^3-83/hCk^ mice. Data were compared using a Mann-Whitney test. **d,** Summary graphic of peripheral B cell development in *Igk*^3-83/hCk^ mice. **e,f,** Variant kappa mice were crossed with MD- and M-only mice. Light chain expression was monitored in the immature, transitional and naïve compartments, with **e,** representative FACS plots and **f,** dot plot showing the percentage of huIgK^+^ and/or IgL^+^ cells. **b,c,f,** Dots represent values from a single mouse. Genotypes were compared pairwise using a Tukey’s post-hoc test, and *P*-values are as marked. **g,h,** BCR sequences from the mature naïve B cell compartments of MD- and M-only mice were obtained, cloned and expressed as IgG. Antibodies were screened against dsDNA, ssDNA, histone, insulin, lipopolysaccharide (LPS), keyhole limpet hemocyanin (KLH) and cardiolipin (CL). ELISA traces show the binding curves antibodies derived from MD- (red) and M-only (blue) mice, and ED38 positive control (yellow) and mGO53 negative control (green)^42^. **h,** Pie charts show the fraction of antibodies that were more reactive than mGO53 against at least 3 different antigens. Data were compared via a right-tailed Fisher’s exact test.

To evaluate how IgD expression impacts peripheral negative selection, we crossed the variant IgK mice to MD- or M-only models, and compared expression of huC_k_ and IgL in the mature naïve compartment. The proportion of huIgK- or IgL-expressing mature naïve B cells was significantly greater in M-only compared with the MD-only background (*P* < 0.0001; Fig. 3e,f). The data suggest that B cells expressing IgD and 3-83_IgK_^+^ are more likely to accumulate in the mature naïve compartment than their IgM expressing counterparts.

To directly determine whether IgD expression is permissive of polyreactive naïve B cell survival, we assessed the polyreactivity of antibodies cloned from the naïve B cells of both MD- and M-only mice by ELISA (Fig. 3g)^42^. The proportion of polyreactive naïve B cell antibodies obtained from MD-only mice was significantly greater than their M-only counterparts: absence of IgD resulted in a 33% reduction in polyreactive clones (Fig. 3g,h; *P* = 0.048). The data are consistent with the idea that expression of IgD increases the threshold for negative selection and thereby supports an expanded naïve B cell compartment^37^.

Under physiologic conditions, IgD expression is extinguished after B cells encounter their cognate antigen, receive T cell help, and enter GCs. Nevertheless, high affinity anti-NP serum antibodies produced by mice that carry both a WT and an IgD deleted allele tend to be produced by B cells expressing the intact Ig allele^18^. One potential explanation for these apparently incongruous results is that IgD sets thresholds for B cell entry into GCs. To determine how IgD expression impacts mature naïve cell selection in response to foreign antigen, we immunized mice with NP-conjugated ovalbumin (NP-OVA) and analysed GCs in draining popliteal lymph nodes (pLNs) shortly after initiation of the response (Fig. 4a). GCs were similar in size (Supplementary data fig. 9a,b), but the fraction of NP-binding cells as measured by flow cytometry using NP-BSA, was significantly greater in MD-only mice compared with their M-only counterparts (*P* = 0.008; Fig. 4b). Similarly, MD-only mice immunized with the diphtheria and tetanus vaccine (Tenivac), or SARS-CoV-2 receptor binding domain (RBD) showed a greater proportion of antigen-binding GC B cells than their M-only counterparts (Fig. 4c,d; Supplementary data fig. 9c–e). Thus, IgD expression results in recruitment of a higher proportion of B cells expressing antigen receptors with demonstrable antigen binding activity, suggesting more stringent selection into the GC. Like their WT counterparts, MD-only B cells in established GCs downregulate IgD (Supplementary data fig. 9f)^45^. Consistent with this change in isotype expression, the proportion of antigen-binding cells in MD- and M-only mice normalized under selective pressure with time in the GC (Fig. 4d; Supplementary data fig. 9g).

**Fig. 4:**
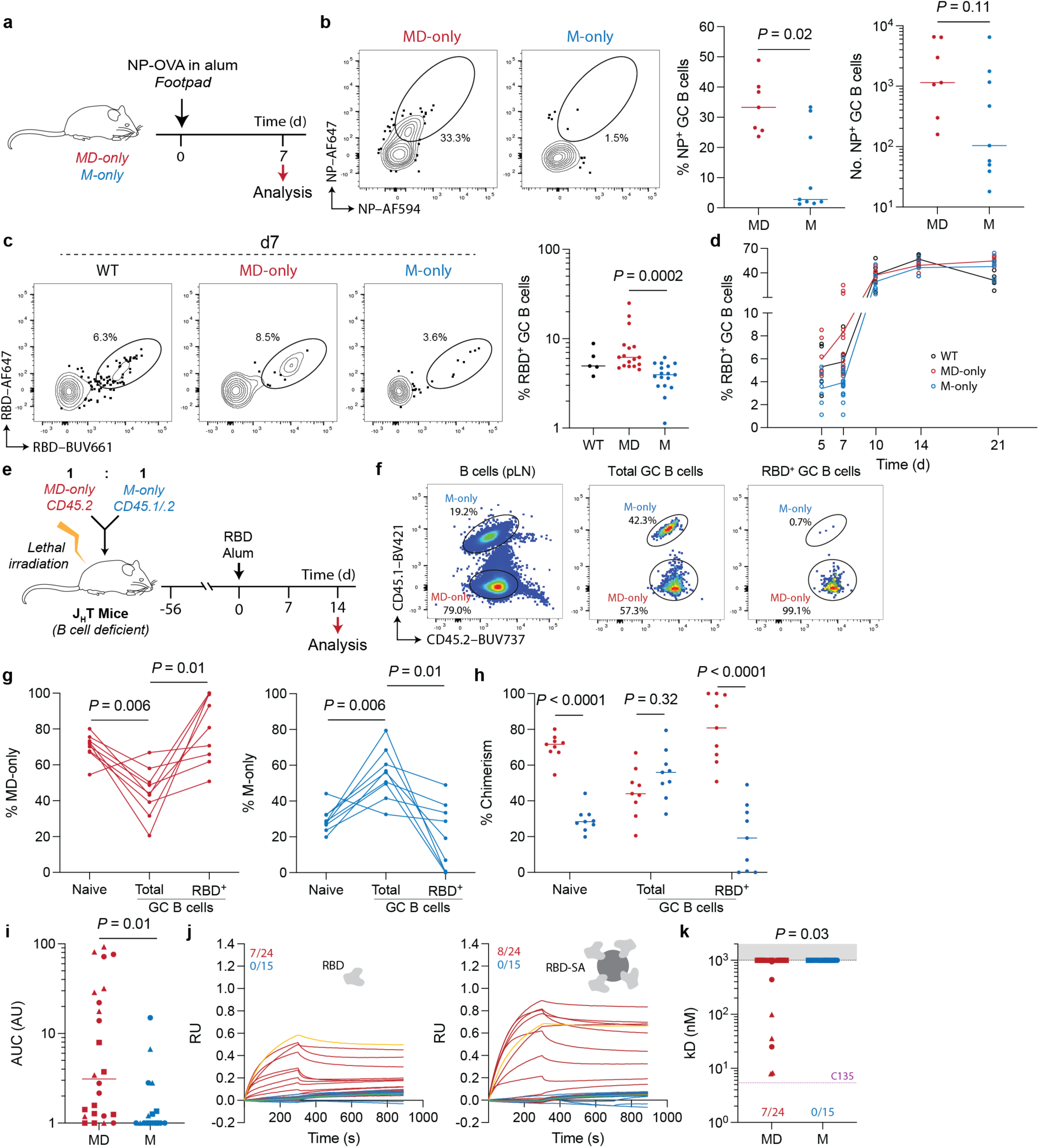
Naïve cell expression of IgD improve selection stringency of clones entering the GC. **a,** Diagrammatic representation of the immunization schedule for b. **b,** Representative flow cytometry plots show the proportion molecular bait-stained GC B cells. Dot plots show the proportion and absolute number of bait-positive GC B cells in each mouse. Groups were compared by a Mann-Whitney test. **c,** Immunization with SARS-CoV-2 RBD. The proportion of RBD-binding GC B cells was evaluated over time by flow cytometry. Lines represent the median for each genotype. Flow cytometry and summary dot plots of d7. **d,** Plot shows the kinetics of RBD bait-binding GC B cells over time. **c,d,** Dots represent individual mice, *n* = 5–18 per group, pooled from at least two independent experiments. Bars represent the median. Data were compared by Kruskal-Wallis test and post-hoc Dunn’s multiple comparison. **e,** Experimental scheme for f–h. J_H_T mice^59^ were reconstituted with 1:1 MD-only (CD45.2) and M-only (CD45.1/.2) bone marrow and immunized with RBD. Germinal centers in the pLN were evaluated after 14 days. **f,** Representative flow plots show the proportion of congenically discordant B cells in different compartments. **g,h,** Graphs show the chimerism of B cells across the compartments. Dots represent data from a single animal, with conjoined lines denoting the same mouse. ANOVA was performed and pairwise comparisons were made via paired repeated measure (RM) test. **i–k,** IgG were cloned from GC B cells on d7 after immunization with RBD. **i,** Plot showing the calculated area under the curve (AUC) as determined by ELISA. **j,** BLI traces of individual IgGs are shown: MD-only-derived IgG (red), M-only-derived (blue), the C135 high-affinity positive control (yellow)^56^, and mGO53 negative control (green)^42^. IgGs were immobilized to the probe, and analytes of either monomeric RBD (left) or tetramized RBD (right) were screened. **k,** Graph shows the k_D_ values of the antibodies tested in j. The limit of detection was k_D_ = 1 µM and the C135 positive control bound at k_D_ = 5.4 nM. Shapes represent clones from different animals (*n* = 3 mice per group). The proportion of detectably reactive clones were compared by a chi-square test.

To confirm that IgD expression raises the threshold for GC entry, we immunized MD-and M-only mixed bone marrow chimeric mice with the SARS-CoV-2 RBD and examined their GCs shortly after initiation which corresponds to 14 days after immunization in our bone marrow chimeras (Fig. 4e–h). Whereas MD-only origin cells were significantly overrepresented in the total B cell compartment compared with M-only cells, they were underrepresented in the GC, suggesting a higher threshold for GC entry (Fig. 4f–h; *P* = 0.007). Notably, however, MD-only B cells were significantly overrepresented among all antigen-binding GC B cells and made up a mean of 80% of these cells (Fig. 4g,h; *P* = 0.01). These data are consistent with the idea that IgD expression increases the threshold for antigen dependent selection.

To measure the relative affinity of antibodies expressed by MD- or M-only B cells recruited into the GC, we obtained Ig sequences from single cells 7 days after immunization with RBD when somatic hypermutation is still low (Supplementary data fig. 9h,i). Binding to RBD was measured by ELISA and bio-layer interferometry (BLI) for 24 MD- and 15 M-only antibodies (Fig. 4i–k; Supplementary data fig. 9j). 83% of antibodies derived from MD-only mice detectably bound antigen in ELISA assays, compared with 50% of M-only antibodies (*P* = 0.03; Supplementary data fig. 9j). As expected, the affinity of IgGs cloned from the early GC was generally low, but MD-only antibodies showed higher affinities than M-only antibodies (Fig. 4i,j). The data confirm that IgD expression is associated with a higher affinity threshold for recruitment into the GC.

The data suggest that the IgD hinge reduces B cell sensitivity to antigen during both negative and positive selection. To determine whether selection thresholds are directly related to the hinge, we deleted the hinge exon from the genome to produced mice with an otherwise intact *Igh* locus but that express a truncated form of IgD whose hinge resembles that of IgM (IgD^trunc^; Fig. 5a–c, Supplementary data fig. 10a and b). Consistent with the findings in M-only mice, IgD^trunc^ mice exhibited lower levels of surface BCR, and a decreased fraction of mature naïve B cells compared with the WT (*P* = 0.001; Fig. 5d; Supplementary Data Fig. 10c). To examine how hinge truncation impacts GC selection, we produced WT:IgD^trunc^ mixed bone marrow chimeras in B cell deficient J_H_T recipients (Fig. 5e). Like M-only mice, the proportion of IgD^trunc^ B cells decreased in the transition between immature and mature B cell compartments relative to the WT (*P* = 0.02; Fig. 5f). To determine how the hinge impacts GC selection, we immunized WT:IgD^trunc^ chimeras with SARS-CoV2 RBD. The relative proportion of IgD^trunc^ GC B cells increased relative to their naïve counterparts while the proportion of WT GC B cells decreased (*P* = 0.006; Fig. 5g). But despite the relative disadvantage of WT GC B cells, they showed a higher proportion of antigen binding (*P* = 0.02; Fig. 5g). Thus, IgD hinge truncation phenocopies the M-only mouse. We conclude that the conserved IgD hinge is essential for raising the thresholds for positive and negative B cell selection.

**Fig. 5:**
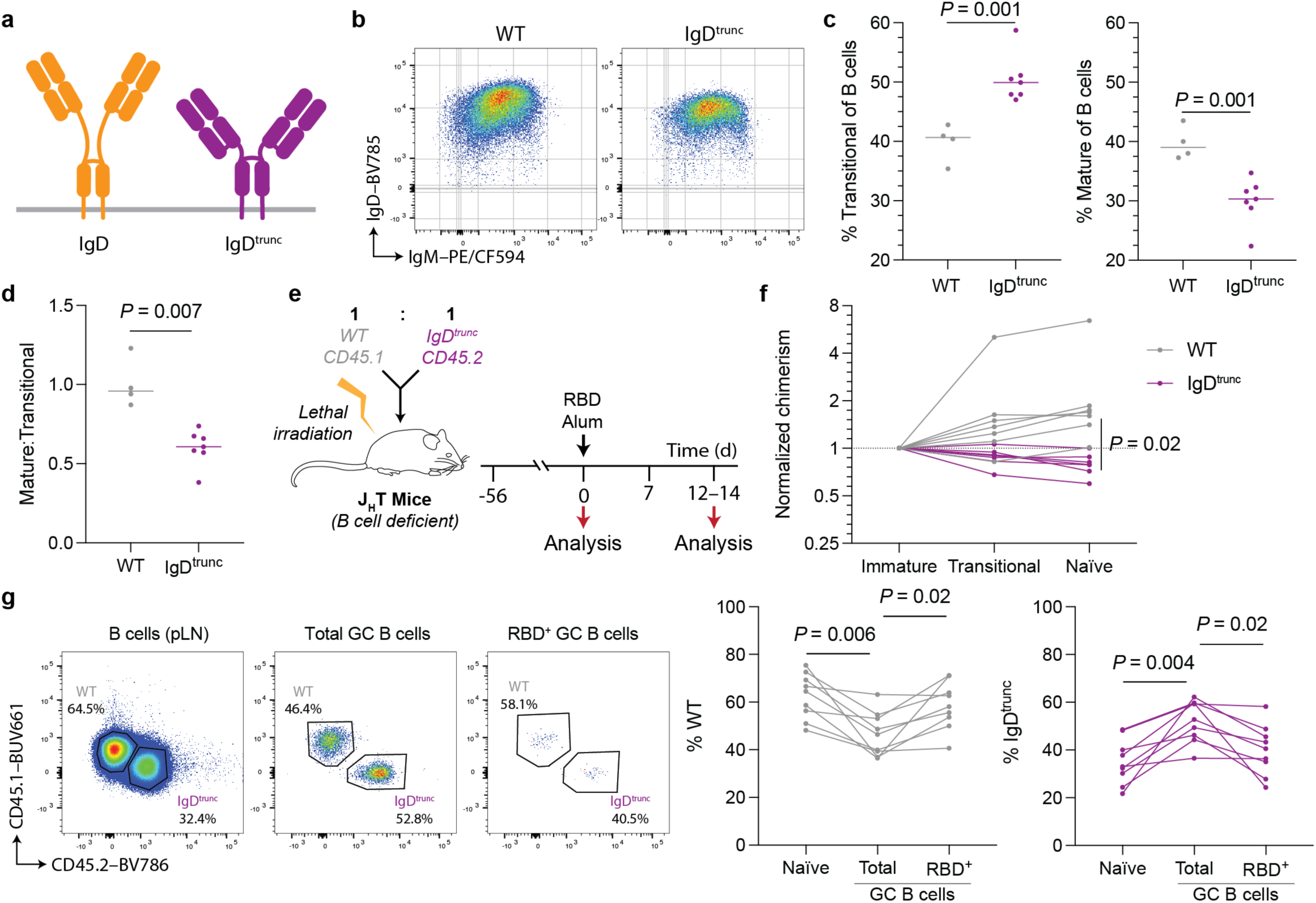
Hinge truncation of IgD perturbs selection thresholds of transitional and naïve B cells. **a,** Graphical depiction of IgD and IgD^trunc^ (Supplementary data fig. 10a). **b,** Representative flow cytometry plots show the expression of both IgM and IgD on mature naïve B cells in WT and IgD^trunc^ mice. **c,** The fraction of transitional (CD93^+^) and mature naïve (CD93^-^CD23^+^) B cells in blood. **d,** Plot shows the transitional-to-naïve cell ratio in blood. **e,** Experimental scheme for f,g. Mixed bone marrow chimeras were generated with lethally irradiated J_H_T hosts, engrafting WT (CD45.1) and IgD^trunc^(CD45.2) marrow. Following reconstitution, animals were immunized with RBD precipitated in alum. **f,** The chimerism of B cells in the periphery (transitional and mature naïve) were enumerated and normalized to the bone marrow immature B cells. **g,** GC B cells were evaluated 12–14 days post-immunization. Representative flow cytometry plots and dot plots show the chimerism among naïve, GC and RBD bait-binding GC B cells. **c,d,f,g,** Dots represent data from a single mouse (*n* = 4–9 per group), with lines conjoining them connoting data from the same animal. Experiments were repeated at least twice *P*-values are as indicated. **c,d,f,** Data were compared using a t-test. **g,** Data were compared using an RM one-way ANOVA and Tukey’s post-hoc multiple comparison.

## Discussion

Despite its strong evolutionary conservation and its unique biophysical features, the function of IgD in physiology has remained enigmatic. Here, we investigated the role of IgD’s unique hinge region in homeostasis and adaptive immunity. We provide a theoretical framework that posits that IgD’s distinct physical properties modulate antigen binding and thresholds for selection. The model is supported by single molecule biophysical measurements and genetic experiments in mice. The data indicates that the IgD hinge minimizes peripheral negative selection and increases naïve B cell activation thresholds thereby optimizing positive selection in response to foreign antigen.

IgD’s biophysical properties have important consequences for peripheral B cell selection and repertoire maintenance. In our transgenic models, the presence of IgD attenuated negative selection in the periphery, enabling a more expansive naïve repertoire that includes B cells expressing polyreactive antibodies. Conversely, despite lower levels of surface BCR, M-only and IgD^trunc^ B cells exhibited enhanced peripheral deletion, consistent with a lower activation threshold. Thus, as suggested by both mouse and human experiments, the canonical IgD BCR serves as a tolerance-promoting element, dampening signaling from low-avidity interactions that might otherwise lead to deletion^12,37,46,47^. Our experiments provide a biophysical explanation for how IgD expression by naïve B cells preserves diversity by favoring anergy as opposed to B cell deletion^46,48^. Moreover, the data indicate that this effect is IgD hinge region dependent. The observation that IgD dampens signaling simply by altering the antigen binding kinetics of the BCR may also explain observed differences between the ability of antigen and anti-BCR crosslinking to elicit intracellular signaling by IgM and IgD^37,38,46^.

Finally, the data reveal that the modulatory effects of IgD extends into the B cell activation phase of the immune response, where it impacts selection of naïve B cells into the GC. By raising the threshold for activation, IgD limits entry of low-affinity clones into the GC, thereby optimizing clonal competition and affinity maturation. This dual role—maintaining an expanded repertoire while calibrating entry into productive immune responses—positions IgD as a determinant of B cell fate. Together, our findings reveal that the evolutionary conservation of IgD, and its long hinge, reflects its unique contribution in balancing both sensitivity and specificity of antigen recognition, promoting both immune tolerance and clonal selection.

## Methods

### Recombinant IgM and IgD

Monomeric *C*-terminally His_6_-tagged native soluble murine IgM (UniProt: P01872) and IgD (P01881) were cloned into mammalian expression vector, pCMV. The cystine-rich tail piece of IgM was removed to prevent oligermerization^49,50^. The corresponding light chain (IgK and IgL) was also cloned into pCMV. Clone details/variable regions as indicated and referenced in the text. Recombinant antibody was expressed by transient transfection in Expi293 cells, as per the manufacturer’s instructions (Life Technologies). Antibodies were first purified from clarified supernatant using immobilized nickel resin affinity chromatography (Life Technologies) followed by up to two rounds of size exclusion chromatography (SEC). Protein was flowed through the Superdex 200 Increase column (Cytiva), collecting fractions validated through both native gel electrophoresis and negative stain electron microscopy to correspond with monomeric Ig. Purified Ig was flash frozen and stored at -80°C.

### Negative stain electron microscopy

Negative staining and imaging were adapted from previously described methods^51,52^. Briefly, glow-discharged copper grids (CF400-CU-50, Electron Microscopy Sciences) were used. 2 x 20 μl drops of distilled water and 2 x 20 μl drops of 1% (w/v) uranyl formate solution were suspended on plastic film. A grid was held with reverse-force anti-capillary forceps, with the carbon-coated side facing upward. Protein sample (2 μl) was applied to the grid and incubated for 30 s to allow adsorption. The grid was then blotted with filter paper, washed twice with water and subsequent blotting, and stained by sequential contact with two drops of 1% uranyl formate: an exposure with the first drop followed by blotting, and a 20 s incubation with the second drop before blotting.

Micrographs were recorded on a JEOL JEM-1400Plus transmission electron microscope operating at 120 keV using SerialEM automated image-acquisition software (http://bio3d.colorado.edu/SerialEM/index.html). Images were acquired at a nominal magnification of 50,000 x, corresponding to a calibrated pixel size of 1.99 Å. Raw images are deposited in Figshare [doi scheduled for release with publication].

### Single-molecule FRET

We engineered recombinant chimeric murine IgM and IgD proteins, substituting the Fc portion for that of mouse IgG1 (P01868). Heavy chains were *C*-terminally His_6_-Avi-tagged. Antibodies were expressed with light chains with a *C*-terminal A4 tag^33^. Proteins were expressed and purified, as described. Antibodies were biotinylated using BirA biotin ligase according to the manufacturer’s instructions (Avidity). To modify the light chains, antibodies were constituted in reaction buffer (50 mM HEPES, 100 mM NaCl_2_, 10 mM MgCl_2_) at 5 µM, along with AcpS, LD555-CoA (Lumidyne Technologies) and LD655-CoA (Lumidyne Technologies) at a 1:2:10:10 molar ratio. The reaction was left to proceed for 2 h at 22°C. After desalting, light chain modification was assessed by spectrophotometric wavelength scanning and gel electrophoresis.

Single-molecule imaging was performed on a custom-built prism-based total internal reflection fluorescence (pTIRF) platform, as described in detail previously^53^. Samples were illuminated using a 6 W 532 nm laser (Quantum Opus) coupled to a 1:100 biotin-PEG:PEG-passivated quartz microfluidic imaging chamber ^54^ through a custom-built quartz prism (Eksma). Fluorescence emission was collected through a Nikon CFI SR Plan Apo IR 60× 1.27 NA water immersion objective, spectrally separated, and imaged onto two sCMOS cameras (Photometrics Kinetix).

Imaging data was acquired using custom software written in C++ and LabVIEW (National Instruments) and processed with SPARTAN 3.9.5^53^ to extract and correct traces for fluorescence baseline, spectral cross-talk (α), relative brightness (γ), and direct acceptor excitation.

Experiments were performed in an imaging buffer containing 1 x PBS (Gibco, pH 7.4) supplemented with an enzymatic oxygen scavenging system (2 mM protocatechuic acid and 50 nM protocatechuate 3,4-dioxygenase)^55^. Data were collected with continuous illumination at the following integration times and irradiances: 2ms, 1530 W/cm^2^; 5ms, 525 W/cm^2^; 10ms, 320 W/cm^2^. For data collected at 2 Hz, we used stroboscopic imaging with a 100 ms integration time and 25 W/cm^2^ irradiance to collect one frame every 500 ms.

For further analysis, traces were selected based on the following criteria: correlation between donor and acceptor, -1.1 to 0.5; signal-to-noise ratio of fluorescence > 10; mean total intensity > 280 photons; mean total intensity within 2 standard deviations; FRET lifetime > 50 frames; background noise < 70. Median correlation between donor and acceptor fluorescence for each molecule (Supplementary data fig. 1) were calculated using a custom MATLAB script by first determining the correlation for each selected molecule prior to photobleaching of either donor or acceptor. Median correlation was then calculated from the distribution of correlation values from each trace.

### Biophysical modelling

Details on the biophysical theory and modelling are outlined in Supplementary document 1.

### Recombinant antigen expression and purification

Antigens were expressed recombinantly using mammalian expression vectors encoding His_6_-Avi-tagged 7MUT gp120^35^, Wu-1 RBD^56^, Tetanus toxin heavy chain *C*-terminal domain (TTHC) or cross-reactive material (CRM)197^57^. Vectors were transiently transfected into EXPI293 cells, and protein was purified from clarified supernatant using nickel resin chromatography, as before. For the generation of antigen baits, antigens were monobiotinylated using BirA biotin ligase, as before.

### BCR transfection and binding studies

Alternative membrane-tethered isoforms of native murine IgM (P01872) and IgD (P01881) were cloned into pCMV as before, and light chain plasmids were identical to that used to express soluble Ig. The variable regions used are indicated in the main text. We also cloned Cd79a/b (Uniprot: P11911 and 15530, separated a self-cleaving P2A peptide) into pCMV. Three vectors—encoding the heavy chain, light chain and Cd79a/b—were co-transfected into HEK293T cells using lipofectamine 2000 (Life Technologies) and incubated at 37°C, 5% CO_2_ for 12–24 h. Cells were harvested using ice cold PBS supplemented with 1% FBS and 2mM EDTA. Following Fc blocking (Human TruStain FcX, BioLegend), surface BCR expression was validated via flow cytometry, using the following antibody cocktail: anti-mouse IgK–Brilliant Violet (BV)421 (clone: 187.1, BD Biosciences), anti-mouse Cd79a–allophycocyanin (APC) (F11-172, BioLegend), anti-mouse IgM–PE/Cyanine-based fluorescence (CF)594 (R6-60.2, BD Biosciences) and anti-mouse IgD–BV785 (11-25c.2a, BioLegend). Data were collected using a BD FACSymphony A5 cytometer and analysed using FlowJo (v10.10.0 for MacOS).

To assess cell antigen capture, fluorescent molecular bait was prepared. Monobiotinylated 7MUT gp120 was tetramerized with phycoerythrin-conjugated streptavidin (SA–PE, BioLegend). NP bait was prepared by constituting BSA (Sigma) in 0.1M NaCO_3_ (pH = 8.0) and subsequently incubated with *N*-hydroxysuccinimide (NHS)-esterified Alexa Fluor (AF)647 (Life Technologies). Following desalting, the fluorescently labelled BSA was incubated with different concentrations of NHS–NP (Setareh Biotech). After quenching excess, unreacted NP with high concentration glycine and subsequent buffer exchanging, the mean NP density was calculated using OD410. After molecular probes were prepared, cells were incubated with 10 µg/mL fluorescently labelled antigen for 25 mins on ice followed by the addition of anti-light chain—anti-mouse IgK–BV421 (187.1, BD Biosciences) or IgL–Pacific Blue (PB; MHL-38, BioLegend)—for an additional 10 mins. Data were acquired by flow cytometry, as described above.

### BLI

BLI assays were performed on the Octet Red instrument (ForteBio) at 30°C with 1 000 prm shaking. Kinetic assays were performed using Octet high precision protein A (ProA) biosensors. Purified IgG was immobilized to the probe and cognate antigen was used for association/disassociation analysis, as detailed in the main text. Each assay step was performed in 1X kinetics buffer (ForteBio). Briefly, the assay settings were as follows: a) 60 s baseline, 2) 200 s probe loading with Ig, 3) 200 s baseline in buffer, 4) 300 s association in analyte and 5) 600 s disassociation period in buffer. Curve fitting was performed using the Octet analysis software (ForteBio) and k_D_ values were calculated according to prior published models^58^.

### Mice

Animals were housed in a specific pathogen-free facility with *ad libidum* chow and water. MD-only (*IgH*^ΔCγ3–⍺/ΔCγ3–⍺^) and D-only (*IgH*^ΔCμ/ΔCμ; ΔCγ3–⍺/ΔCγ3–⍺^) mice were generated from wild-type C57BL/6J mice (Jackson Laboratories), electroporating embryos with two CRISPR/Cas9 ribonucleoproteins (RNPs) to excise target *Igh* genomic regions (NCBI genomic reference: NC_000078.7 [113222388..115973574]). MD-only mice were generated using a 5’ guide (CTAGGACAAGGATACCCCTG) situated ∼900bp downstream of the C_δ_ 3’ UTR, and 3’ guide (TTACTAGGCTCCTCCATATG) downstream of exon 4 of C_ɑ_. D-only mice were generated by second-round targeting of MD-only mice. C_μ_ and σδ was excised using a 5’ guide (GTACTGGATGATCTGGTGTA) ∼300 bp downstream of the μ enhancer critical to both IgM and IgD expression, and 3’ guide (ATCTACACAGATCCCTCCCA) ∼100 bp downstream of the σδ region. IgD^trunc^ (*Igh* ^ΔCδex2/ΔCδex2^) mice were engineered by excising exon 2 of ΔC_δ_, a symmetrical exon that encodes the discrete 35 a.a. hinge of canonical murine IgD: 5’ AACAGTGCAAGGATGCAGCT guide, and 3’ guide GCAAGCCAGGCCTTATATCC. Mice generated in this study were backcrossed for at least two generations. Mice expressing IgH_A_ (*Igh*^a^, stock: 001317) and CD45.1 (SJL; B6.CD45.1/J, stock: 033076) were purchased from the Jackson Laboratory. M-only (*IgH*^Δδ–α/Δδ–α^) mice were as in Schaefer-Babajew et al.^36^, J_H_T mice as in Gu et al.^59^, hCk mice as in Casellas et al.^44^ and 3-83 mice as in Russell et al.^60^. Six-week-old sex-matched mice were used for all experiments, unless otherwise indicated.

### BCR Internalization assay

Primary B cells were isolated from spleen and enriched using the EasySep mouse B cell isolation kit (StemCell Technologies). Cells were stimulated with 10 µg/mL anti-mouse IgK F(ab’)2 (Jackson ImmunoResearch) and maintained in culture at 37°C. At indicated time points, aliquots of cells were placed on ice to arrest BCR internalization. Cells were stained with anti-mouse B220–BV605 (RA3-6B2, BD Biosciences), anti-mouse IgK–BV421 (187.1, BD Biosciences), and anti-mouse IgL–BUV661 (R26-46, BD Biosciences). IgL^+^ B cells were excluded from the analysis.

### Calcium flux assay

Primary B cells were stained with Indo-1-AM (Life Technologies) according to the manufacturer’s instructions. Cells were stimulated with 10 µg/mL anti-mouse IgK F(ab’)2 (Jackson ImmunoResearch) or 1 µg/mL ionomycin (Sigma). Calcium flux was measured longitudinally by monitoring the UV-violet and UV-blue emission ratio using a BD FACSymphony A5 cytometer.

### Treatments and immunizations

Specific immunization and treatment protocols are outlined in the main text. Briefly, mice were immunized with protein precipitated in 1% Alhydrogel (Invivogen). Tenivac (Sanofi Pasteur) was purchased commercially, where each mouse received 1/20th of a human dose. Formulations were administered into the footpad under anaesthesia.

### Bone marrow chimeras

Sex-matched J_H_T mice of 6–10 weeks-of-age were sub-lethally irradiated (2 x 3 Gy) Donor bone marrow cells derived from 5-week-old female mice were mixed, and a total of 2 x 10^6^ cells were transferred to the recipient intravenously. Animals were left for constitution for at least 8-weeks.

### Flow cytometry and cell sorting

Accucheck counting beads (Life Technologies) were added to single-cell suspensions before direct staining with fluorophore-conjugated antibodies against relevant surface lineage markers. The non-B cell dump panel constituted of AF700-conjugated anti-mouse CD3 (17A2, BioLegend), CD4 (RM4-4, BioLegend), CD8a (53-6.7, BioLegend), F4-80 (BM8, BioLegend) and Gr-1 (RB6-8C5, BioLegend). Other lineage markers are outlined in the figures. Antigen-specific B cell staining was performed using tetramerized monobiotinylated antigen, described earlier. Tetramers were generated using SA-PE, SA-BV605, or SA-AF647. Nitrophenylated and fluorescently labelled BSA was described previously. Staining was performed in darkness on ice for 40 mins. Flow cytometric data were acquired on the BD FACSymphony A5 analyzer. Cell sorting was conducted using the BD FACSymphony S6.

### Single-cell mRNA sequencing

Single cells were sorted into 3 µL of lysis buffer containing PEG-8000, Triton X-100, dNTPs, Oligo dT30VN, and RNase Inhibitor in a 384-well plate. First-strand cDNA synthesis was done using Maxima H Minus Reverse Transcriptase (Life Technologies) according to the manufacturer’s protocol using a custom template-switch oligo in a 2 µM final concentration. cDNA was amplified for 23 cycles using KAPA HiFi HotStart ReadyMix (Kapa Biosystems). Amplicons were purified using aMPure XP beads (Beckman Coulter) at a 0.6X ratio. Full-length cDNA was tagmented using Tn5 (Illumina) for 10 minutes at 55°C and libraries were prepared with the Nextera XT DNA Library Preparation Kit (Illumina) with Phusion High-Fidelity DNA Polymerase (Life Technologies) in a total volume of 6.25 µL with 12 cycles of PCR. Indexed libraries were pooled by volume according to plate quadrant and cleaned by aMPure XP beads (Beckman Coulter) at a 0.8X ratio. Pools were sequenced on a NovaSeq X in a PE100 run using the NovaSeq X 10B or 25B Reagent Kit (Illumina). An average of 41 million paired reads were generated per plate.

For targeted BCR sequencing on singly sorted B cells, we performed cDNA synthesis followed by semi-nested PCRs to amplify heavy and light chain sequences. These amplicons were sequenced via Sanger. The tissue processing, cell sorting and reaction conditions were performed as described previously^61^.

### Single-cell library processing

SmartSeq3 libraries reads were trimmed using cutadapt v5.1^62^, and aligned to the mouse GRCm38/mm10 genome build using STAR v2.7.11b^63^ with default parameters except for ‘--limitSjdbInsertNsj 2000000 --outFilterIntronMotifs – RemoveNoncanonicalUnannotated’. Reads were assigned with FeatureCounts^64^, UMI reads were identified with the pattern ATTGCGCAATG^65^, extracted and quantified with umi_tools^66^. The reads were subsequently analysed in R with Seurat v5.1.0^67^. Cells with a mitochondrial content >10% or feature count <200 or >2,500 were discarded. Sample batches were merged, normalized, and scaled with SCTransform. Based on their gene expression profile, single cells were visualized in a lower dimensional space using Uniform Manifold Approximation and Projection (UMAP) clustering. B-cell survival, BCR signalling, and ER stress signatures were obtained from the Molecular signatures database (MSigDB)^68–70^ M2: curated, M5: ontology, and MH: mouse-ortholog hallmark gene sets, and assigned with the Seurat function AddModuleScore. BCR sequences were assembled and analysed with TRUST4 v1.1.7^71^. Contigs containing less than 50 reads and more than one heavy or light chain were removed.

### Bulk BCR repertoire sequencing

Mature naïve B cells (B220^+^CD23^+^CD93^-^) were bulk sorted from splenic single-cell suspensions. RNA was purified using the Monarch Spin RNA Isolation Kit (NEB) according to the manufacturer’s instructions. The NEBNext Immune Sequencing Kit (mouse; NEB) was used to amplify full-length heavy and light chain BCR sequences, using the protocol recommended by the manufacturer. The input total RNA used was 50 ng per reaction. Following PCR purification, samples were screened using the 4200 TapeStation (Aligent) to confirm sufficient amplicon purify. Sequencing was performed on the MiSeq i100 (Illumina).

### Computational analyses of antibody sequences

Processing of bulk BCR repertoire sequencing reads was performed using the publicly available Galaxy workflow: https://usegalaxy.org/u/bradlanghorst/w/presto-nebnext-immune-seq-workflow-v320. This workflow conducts data processing and read alignment of heavy and light chain BCR sequences.

To analyze paired single-cell heavy and light chain sequences acquired via both NGS and Sanger, we used IgPipeline v.3.0 using the default murine immune gene segment reference^72^. Scripts for sequence annotation, processing and graphics rendering are publicly available on GitHub (https://github.com/stratust/igpipeline/tree/igpipeline3).

### IgG cloning, expression and purification

Heavy/light chain variable region gene fragments were synthesized (IDT) and cloned into linearized human IgG1 expression vectors using Gibson assembly as described previously^73^. Recombinant proteins were expressed by transient transfection in Expi293 cells. IgG1 was purified using Protein G agarose resin (Life Technologies).

### ELISA

To evaluate the isotype composition of serum samples, high-protein binding plates were coated with goat anti-mouse Ig antibodies from the C57BL/7 SBA Clonotyping system (SouthernBiotech) overnight at 4°C. Following washing with PBS supplemented with 0.05% Tween-20 (PBST), plates were blocked with 2% (w/v) BSA-PBST. Serum samples were assayed at a starting concentration of 1:100 and serially diluted 3-fold. To determine the absolute concentration of antibodies in the serum, we also assayed immunoglobulin standards from the C57BL/6 mouse immunoglobulin panel (SouthernBiotech). Mouse antibody was detected using horseradish peroxidase (HRP)-conjugated anti-mouse IgK (187.1, SouthernBiotech) and anti-mouse IgL (36-59, Life Technologies). Plates were developed using 1-step Ultra 3,3’,5,5’-Tetramethylbenzidine (TMB; Life Technologies) for 4 mins, and the reaction was arrested by the addition of 0.5 M H_2_SO_4_. Data were acquired using the FLUOstar Omega (BMG Labtech), processed using the MARS analysis software (v.3.20 for Windows; BMG Labtech), and graphics rendered using Prism 10 (v.10.6.1 for MacOS; GraphPad).

Reactivity against self-antigens were conducted as described previously^42^. Briefly, plates were coated with dsDNA purified from Salmon sperm (Life Technologies), ssDNA (prepared by denaturing dsDNA at 95°C for 30 mins), insulin (Sigma), keyhole limpet hemocyanin (KLH; Sigma), recombinant histone octamer (EpiCypher), lipopolysaccharide (LPS) from *Salmonella enterica* (Sigma) or cardiolipin (CL; Avanti Research). For CL immobilization, the lipid was solubilized in anhydrous ethanol, dispensed in the ELISA plate and left at room temperature overnight to allow ethanol evaporation. Wash steps were performed with 0.001% Tween-20. Recombinant antibodies were assayed from a starting concentration of 30 µg/mL and serially diluted 4-fold. Antigen-specific binding of recombinant Fabs was performed, assaying from a starting concentration of 50 µg/mL. Detection of recombinant human IgG1 was performed using anti-human IgG–HRP (H + L; Jackson ImmunoResearch). Development and data acquisition was performed, as above.

### Data processing and statistics

All statistical tests were calculating in Prism 10 or R (v.4.4.1). Specific statistical test details, including *n* and statistical significance values, are indicated in the text and figure legends. For log-transformed data, the geometric mean was used to indicate central tendency, unless otherwise indicated; correspondingly, for non-rank-based statistical tests, groups were compared using the log-transformed data. All comparisons are two-tailed and multiple comparisons are adjusted for false discovery.

### Ethics statement

All animal procedures were performed in accordance with the protocols approved by the Rockefeller University Institutional Animal Care and Use Committee (IACUC).

## Data availability

All data needed to evaluate the conclusions in the paper are available in the main text and supplementary materials. Sequencing datasets have been deposited in the National Center for Biotechnology Information (NCBI)’s Gene Expression Ombibus (GEO) and are available through accession numbers [deposition scheduled upon acceptance]. The raw images acquired for negative stain electron microscopy are deposited in Figshare [doi scheduled for release upon acceptance]. Single-molecule FRET data are available upon request to S.C.B. Source data are provided with this paper. Reagents, including mouse strains, are available from M.C.N. upon reasonable request, under a material transfer agreement with The Rockefeller University.

## Code availability

Custom code generated by the authors are publicly available. The latest version of the IgPipeline software is found at: https://github.com/stratust/igpipeline/. Single-molecule FRET data analysis software, SPARTAN 3.9.5, can be retrieved at: https://github.com/stjude-smc/SPARTAN.

## Acknowledgments

We thank members of the Nussenzweig lab for discussions, T. Eisenreich for animal husbandry, K.-H. Yao for technical support, L. Urnavicius for assistance with negative staining, The Rockefeller University Transgenic and Reproductive Technology Center for performing embryonic electroporation, and K. Gordon for flow cytometry support. We acknowledge the use of the Integrated Genomics Operation Core, funded by the NCI Cancer Center Support Grant (CCSG, P30 CA08748), Cycle for Survival, and the Marie-Josée and Henry R. Kravis Center for Molecular Oncology. We thank D. Terry and the Single-molecule Imaging Center at St. Jude for assistance with single-molecule imaging experiments.

## Funding

This work was supported in part by National Institutes of Health (NIH) grant 5R37 AI037526, NIH Center for HIV/AIDS Vaccine Immunology and Immunogen Discovery (CHAVID) 1UM1AI144462-01 to M.C.N., NIH P01 grant 2P01AI100148-11 to M.C.N, the Stavros Niarchos Foundation Institute for Global Infectious Disease Research, and the St. Jude Children’s Research Hospital Collaborative Research Consortium on G protein-coupled receptors (GPCR). A.J.M. is a Cancer Research Institute Carson Family Fellow. A.K.C. is supported by the Ragon Institute of Mass General Bringham, MIT and Havard University. M.C.N. is a Howard Hughes Medical Institute (HHMI) investigator. L.P.D. is an HHMI Fellow of the Life Sciences Research Foundation.

## Author information

Conceptualization of project: LPD, MCN

Methodology: LPD, RAB, AG, GSS, SZ, HH, DS-B, ZK, TYO, AKC, SCB, MCN

Investigation: LPD, RAB, AG, GSS, SZ, HH, AJM, CU, LB, MT, DS-B, BH, ZK, AG, TYO

Funding acquisition: AKC, SCB, MCN

Project administration: LPD, MCN

Supervision: AKC, SCB, MCN

Writing – original draft: LPD, MCN

Writing – review & editing: all authors

## Competing interests

MCN is on the scientific advisory board of Celldex Therapeutics. SCB has equity interests in Lumidyne Technologies. AKC is a consultant for Flagship Pioneering and its affiliated companies, Apriori Bio and Metaphore Bio; he holds equity in these companies and in Dewpoint Therapeutics. All other authors decare they have no competing interests.

---

**Supplementary data fig. 1.**
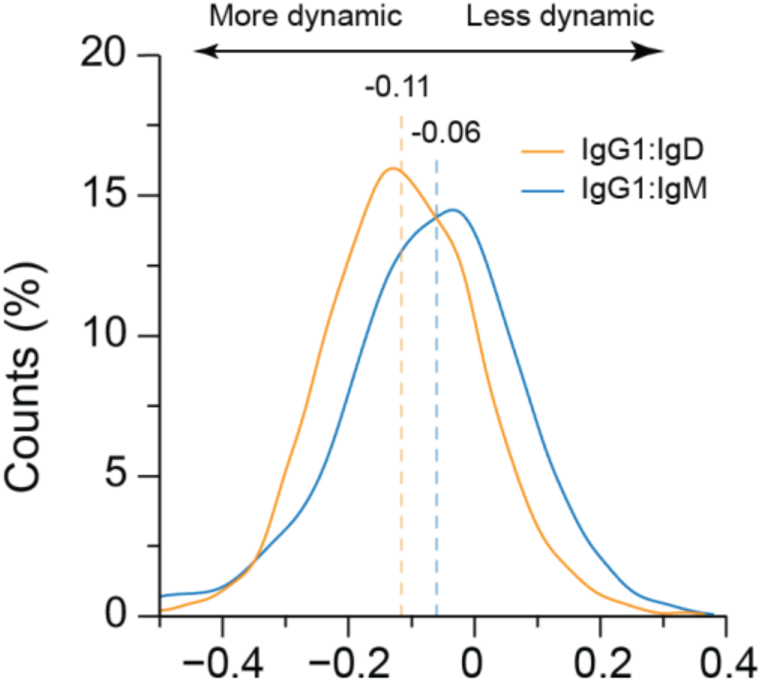
Fab arm dynamics of IgM and IgD hinges measured smFRET. Histograms of Pearson’s correlation of donor and acceptor intensities for IgG1:IgD (4332 molecules) and IgG1:IgM (4291 molecules) from data collected at 10 ms time resolution. Dotted lines denote the median.

**Supplementary data fig. 2.**
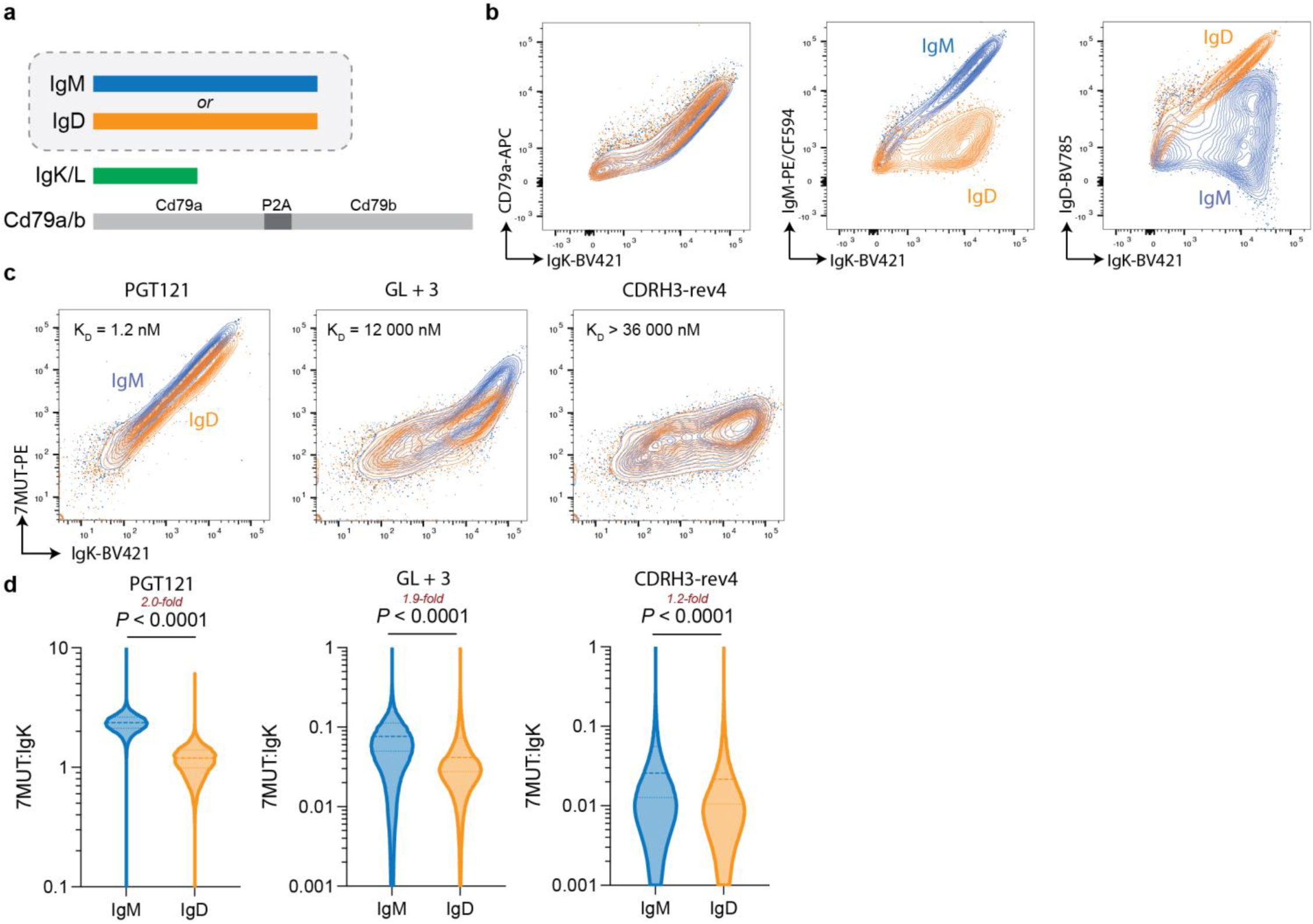
Differential avidity effects of IgD and IgM in antigen capture. **a,** Depiction of the three-plasmid BCR transfection system comprised of plasmids encoding the murine light -chain, Cd79a/b, and either native membrane-bound IgM or IgD. The clone used in each panel are defined. HEK293 T cells were transiently transfected. **b,** Cells were transfected to express the gp120 (7MUT)-binding clone, PGT121, expressed as IgM or IgD BCR. Expression of the respective BCRs were confirmed by flow cytometry. **c,d,** BCR expression of PGT121 and their low-affinity partial inferred germline revertants, GL+3 and CDRH3-rev4 ^(33)^, was induced by transient transfection. **c,** Representative flow plots and **d,** violin plots show the relative dextramerized 7MUT 20-mer capture by cells with respect to isotype. Data were compared using a lognormal t-test.

**Supplementary data fig. 3.**
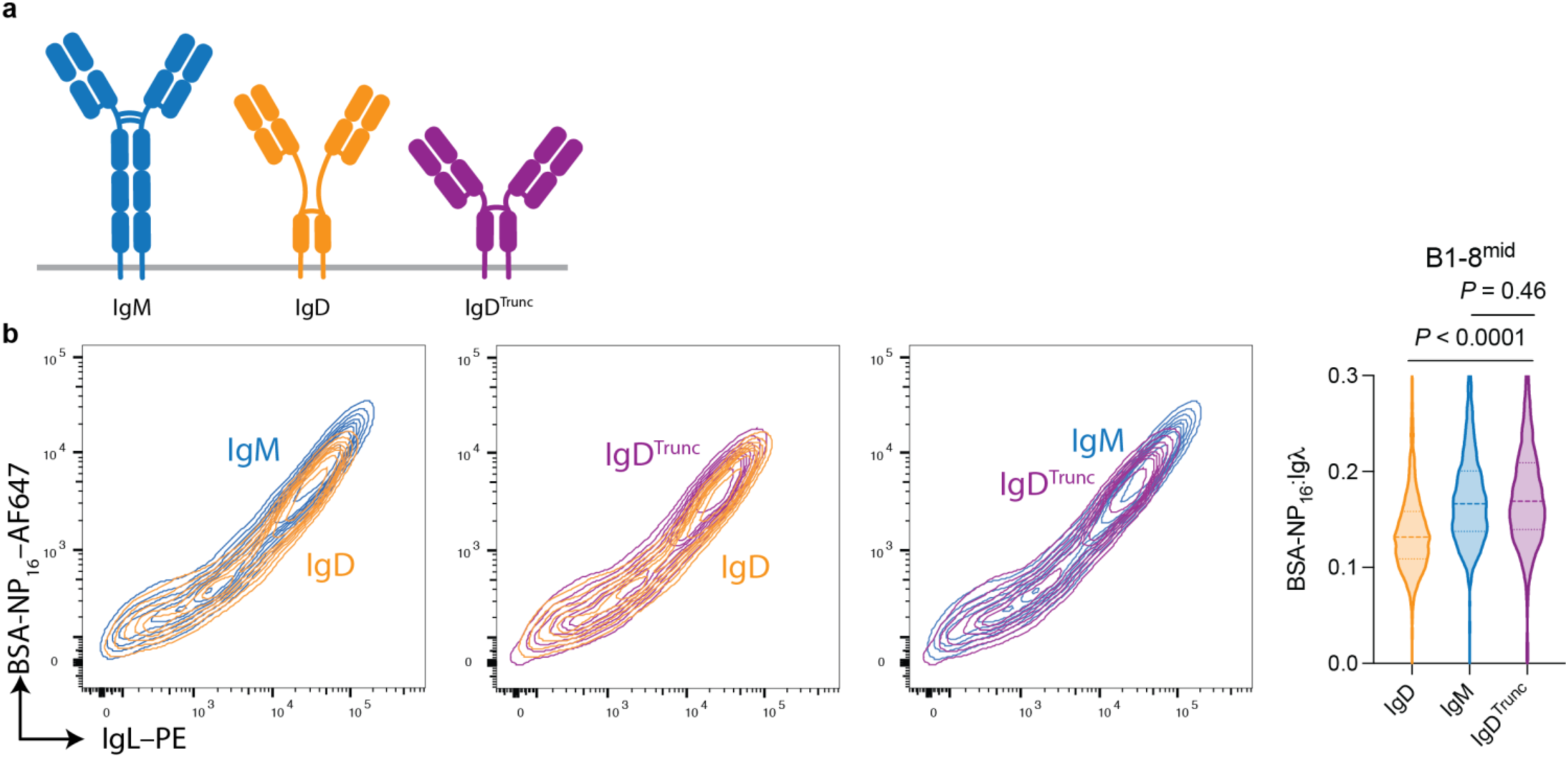
Truncation of the IgD hinge rescues BCR antigen capture. **a,** Graphical depiction of IgD^Trunc^ alongside native murine IgD and IgM BCR. IgD^Trunc^ was engineered to remove the 35 a.a. hinge of IgD (Δ94–128 a.a.; Uniprot: P01881). **b,** HEK293T cells were transiently transfected to express the NP-binding antibody B1-8^mid^ as a BCR (IgH, IgL and Cd79a/b). Antigen capture was quantified by flow cytometry using fluorescently-labelled BSA-NP_16_ and normalizing to the surface BCR density. Representative flow plots show comparative antigen binding with respect to heavy chain, and violin plots show the ratio of antigen- and IgL-associated fluorescence. *P*-values were determined by Tukey’s post-hoc comparison.

**Supplementary data fig. 4.**
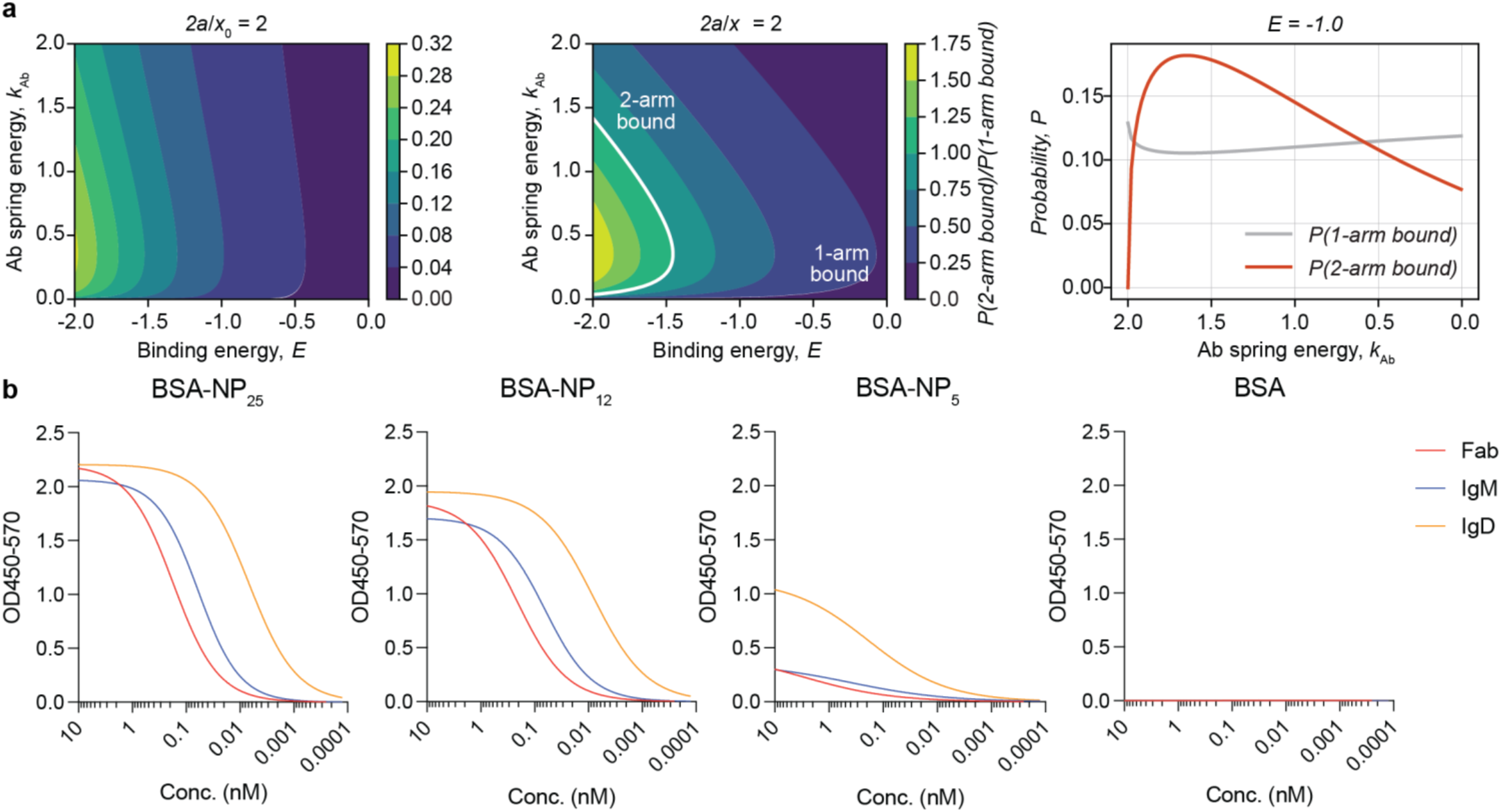
Differential binding of IgM and IgD against fixed antigen. **a,** Plots show the probability of Ig binding to immobilized antigen that deviates from the preferred topology (2a/x _0_ ≠ 1), specifically where epitopes are more distant (2a/x_0_ > 1). **b,** Plots show the ELISA traces of soluble B1-8^hi^ expressed as Fab, murine IgM and IgD binding to BSA-NP_25_, or BSA-NP_12_, or BSA-NP_5_, or BSA. Traces reflect the averaged logistic regression function (*n* = 2).

**Supplementary data fig. 5:**
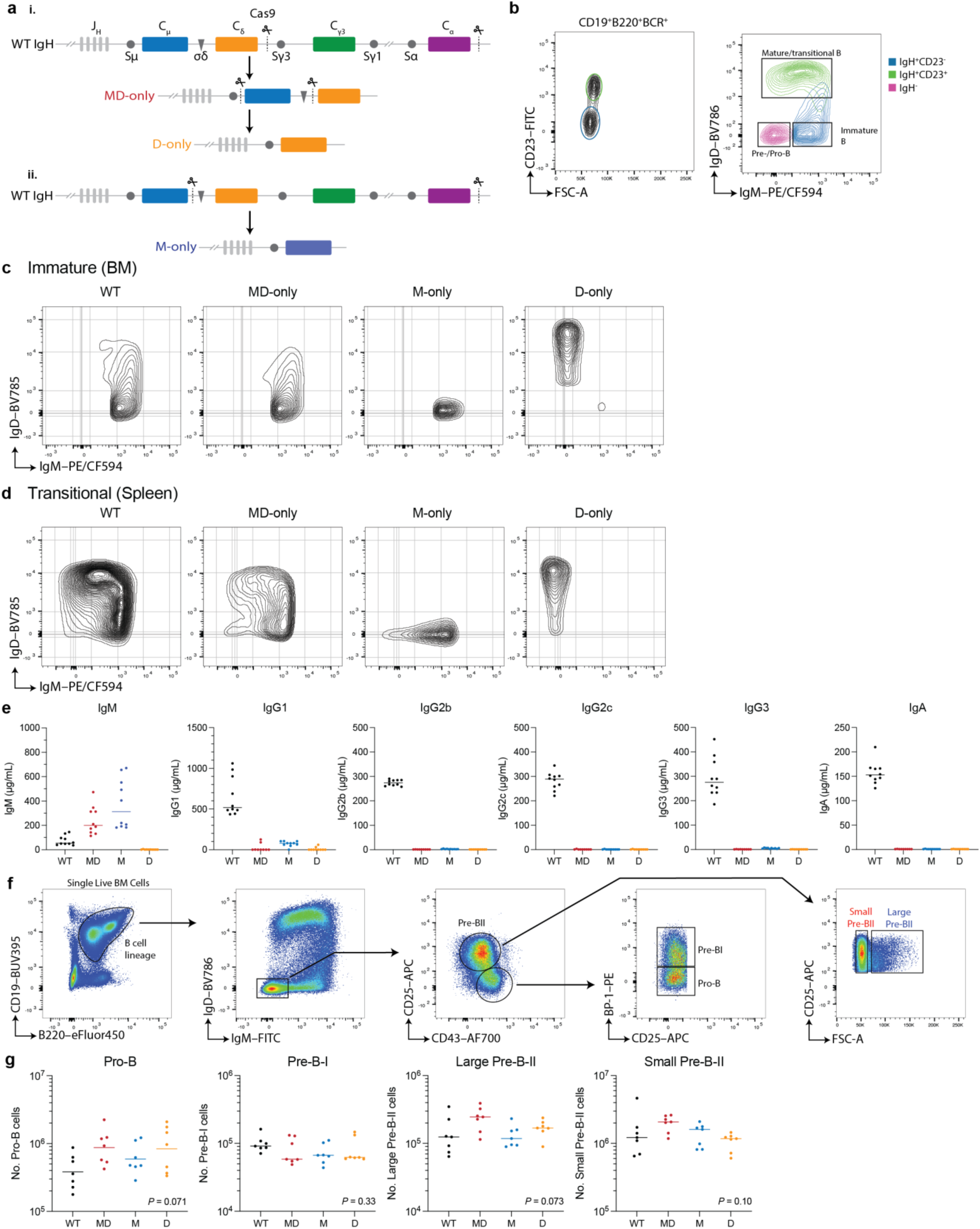
Immune phenotyping of MD- M- and D-only mice. **a,** Graphical depiction of the *Igh* loci and Cas9 gene segment excision strategy to produce i) MD-only mice and subsequently D-only mice by a second round of targeting, and ii) M-only mice. See methods section. **b,** Representative flow cytometry plots of wild-type bone marrow B cells show that B220^+^BCR^+^CD23^-^ immature B cells (IgM^mid–hi^IgD^-^). Flow cytometry plots show the IgM and IgD expression profiles of **c,** bone marrow immature (BCR^+^CD23^-^) and **d,** splenic transitional (CD93^+^) B cells from WT, MD-only, M-only and D-only mice. **e,** Plots show the serum concentration of Ig isotypes as determined by ELISA. **f,** Representative flow cytometry plots show the gating strategy used to identify early B cell precursors in the bone marrow. **g,** Dot plots show the number of B cell precursors per femur. Data were compared by one-way ANOVA. *P* values are marked. **e,g,** Dots represent data from a single mouse (*n* = 7–10 per group) and bars represent the median. Experiments were repeated at least twice.

**Supplementary data fig. 6:**
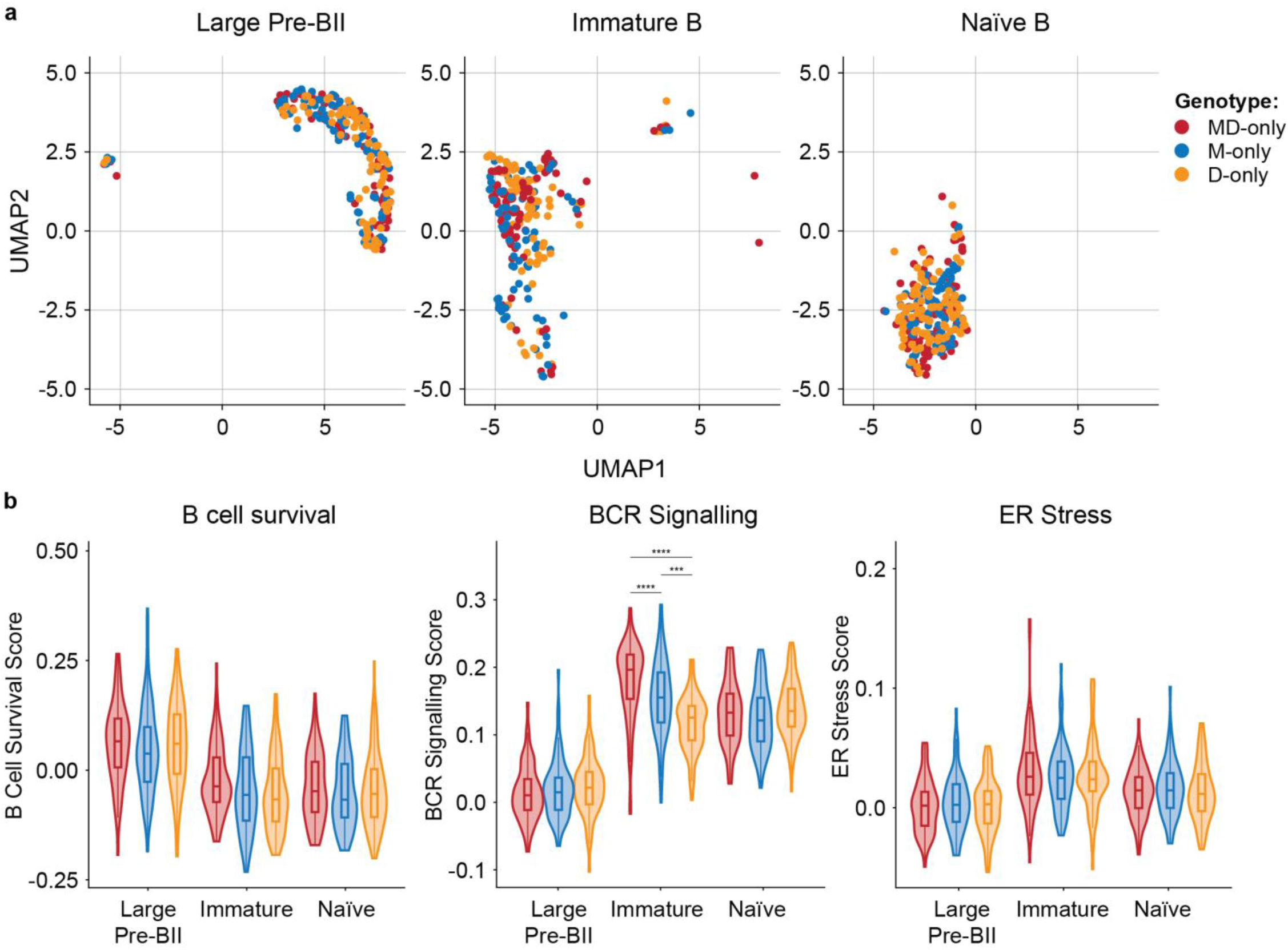
Single-cell RNA sequencing of B cell compartments. Single-cell RNA sequencing was performed on large pre-BII, immature and mature naïve B cells isolated from MD-, M-and D-only mice. **a,** UMAP plots depicting large pre-BII, immature and naïve B cell clusters. Dots represent data from a single cell, and the color denotes the genotype and B cell stage of the corresponding cell. **b,** Violin plots show gene expression pathway scores. Central boxes denote the quartile bounds. Data were compared via Mann-Whitney tests with Benjamini-Hochberg adjustment. *P*-value denotations: ‘***’ *P* < 0.001 and ‘****’ *P* < 0.0001.

**Supplementary data fig. 7:**
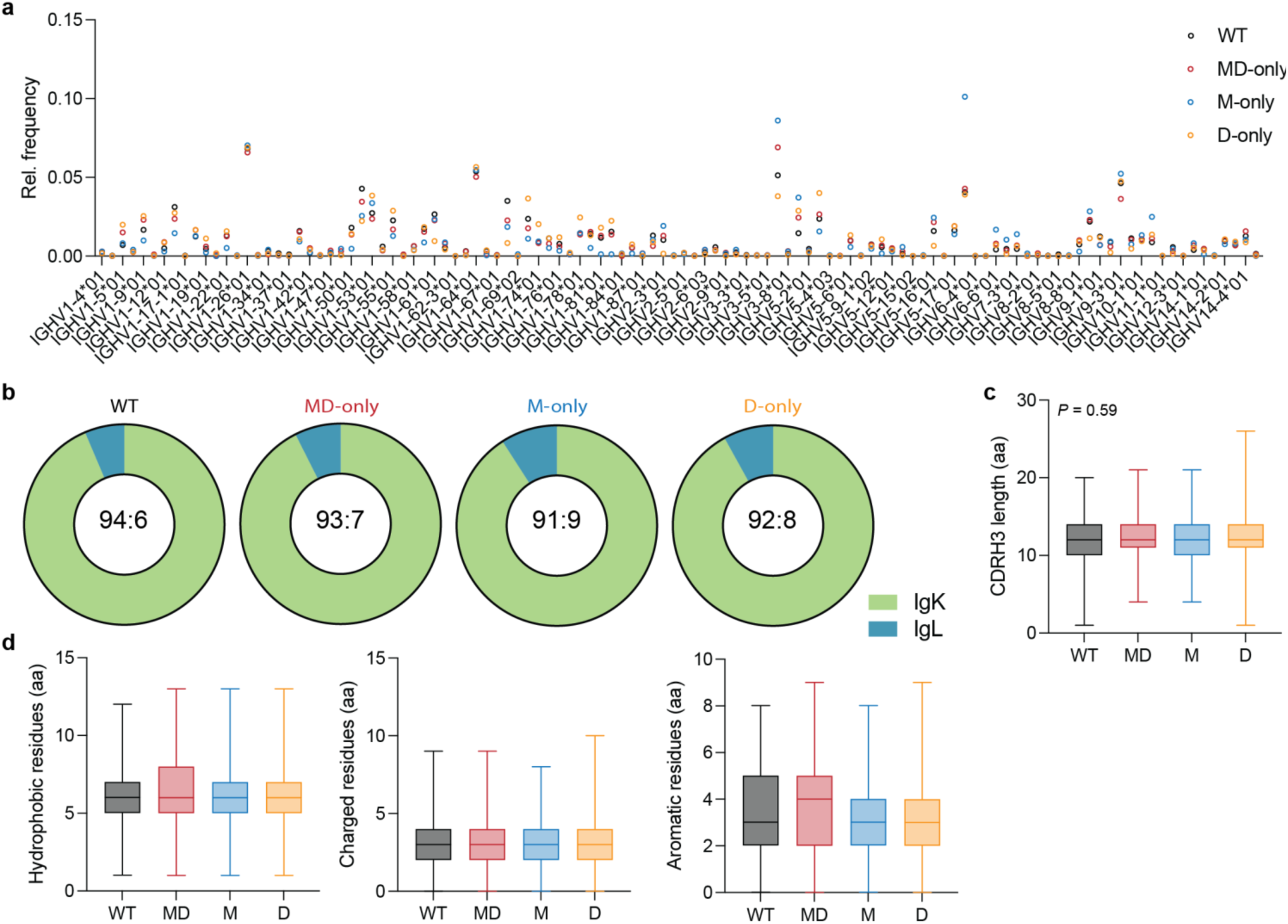
Naive repertiore analysis of MD-, M- and D-only mice. Variable regions from mature naïve (CD23^+^CD93^-^CD43^-^) B cells were sequenced from MD-, M- and D-only mice (3 mice pooled per group). **a,** Plot shows the relative frequency of *Ighv* gene segments for each genotype. **b,** Pie charts show the ratio of IgK and IgL transcript expression. **c,** Box plot shows the quartiles of CDRH3 lengths of naïve B cells with respect to genotype and **d,** shows the number of hydrophobic, charged and aromatic residues. Data were compared by one-way ANOVA.

**Supplementary data fig. 8:**
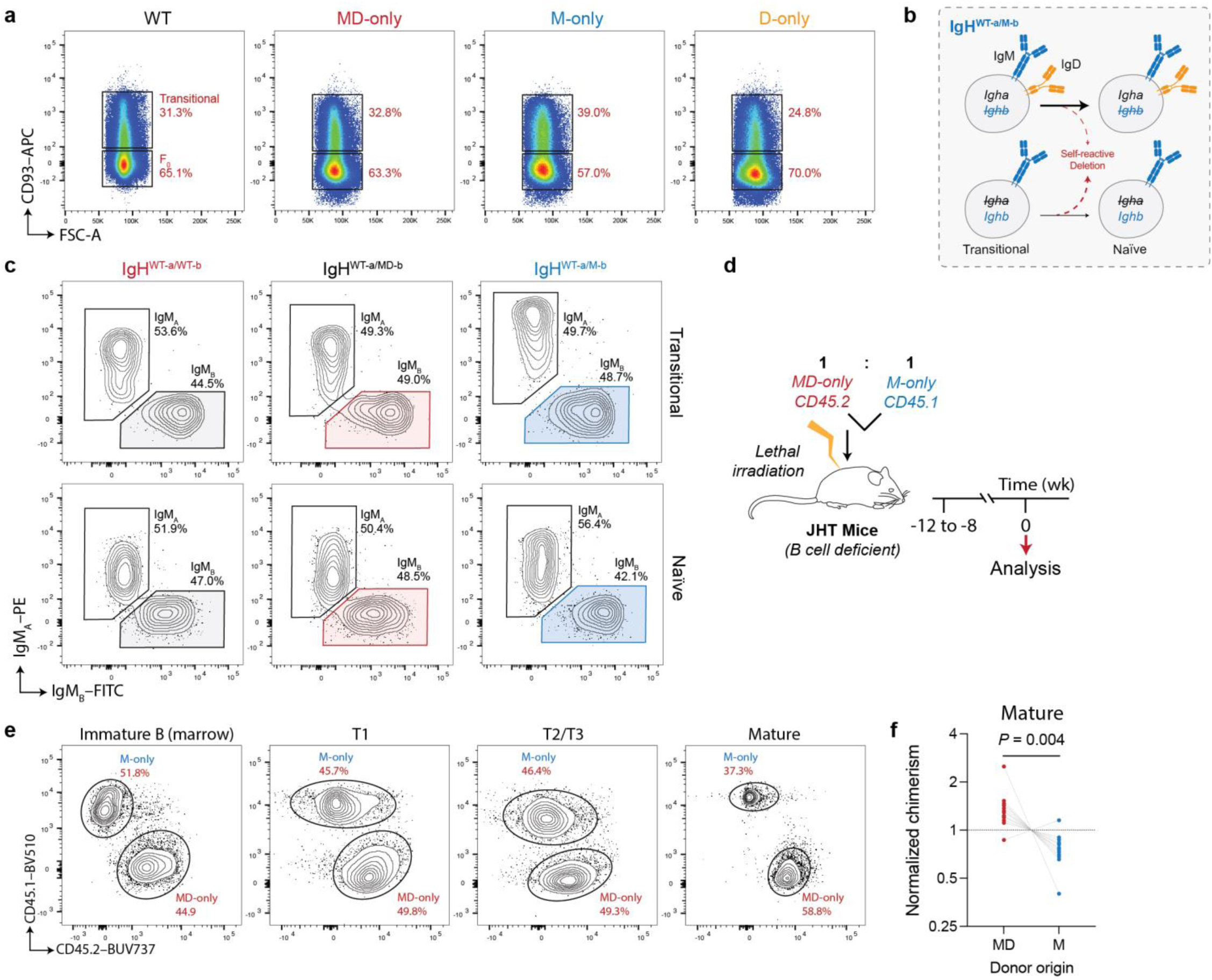
IgD expression in peripheral B cell development. **a,** Representative FACS plots showing the proportion of splenic transitional and mature naïve B cells. Data were pre-gated on DUMP^-^B220^+^CD43^-^ events. **b,** Diagrammatic representation of B cell peripheral deletion in IgH^WT-a/M-b^ mice. **c,** Representative contour plots of allotype expression in the transitional and mature naïve B cell compartments of discordant IgH^WT-a/WT-b^, IgH^WT-a/MD-b^ and IgH^WT-a/M-b^ mice. **d,** Diagrammatic representation of chimera production. 6–8-week-old J_H_T mice ^(48)^ were lethally irradiated and bone marrow obtained from MD-only.CD45.2 and M-only.CD45.1 mixed (1:1) was administered intravenously. After 8–12 weeks, the reconstitution of B cells was evaluated by flow cytometry. **e,** Representative contour plots showing donor B cell origin, as inferred from CD45.1 or CD45.2 expression. **f,** Plots show the relative chimerism, normalized to the immature B cell compartment. Joined dots represent data from the same mouse. This experiment was repeated three independent times. Data were compared by paired t-tests.

**Supplementary data fig. 9:**
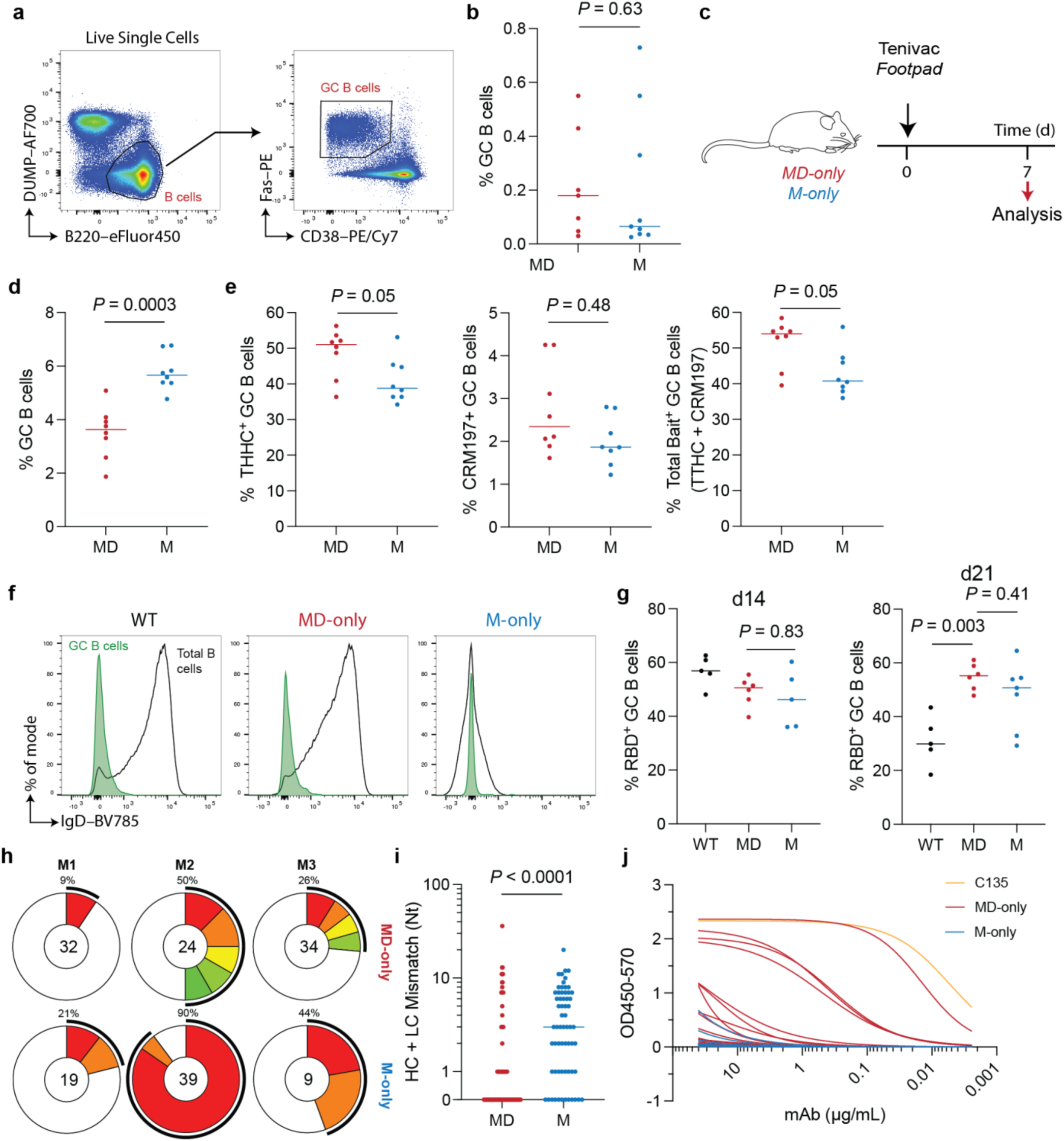
Clonal seeding and early participation of B cells in the germinal center. **a,** Plots show the gating strategy for GC B cells in the pLN. **b**, Graphs show the percentage of GC B cells in mice immunized 7 days prior with NP_17_-OVA. **c,** Immunization schedule for panels (D and E). MD- and M-only mice were immunized with Tenivac and their germinal centers were evaluated after 7 days. **d,** Plot shows the germinal center size in mice immunized with Tenivac. **e,** Plots show the percentage of Tenivac antigens THHC- and CRM197-bait-binding GC B cells post-immunization. **f,g,** Mice were immunized with RBD in alum, and fluorescently-labelled RBD bait staining of GC B cells was quantified longitudinally by flow cytometry. **f,** Representative histogram plots show the IgD expression on GC B cells 14 days post-immunization. **g,** Dot plots show the percentage of GC B cells that bound RBD molecular bait 14 and 21 days after immunization. **h,** Donut plots show the clonal distribution of paired Ig sequences (IgH and IgK/IgL) of GC B cells from MD- and M-only mice immunized 7 days prior with SARS-CoV2 RBD. Colored segments represent the proportion of cells isolated from the same inferred clonal family, whereas the white segment represents singlets. The number inside of the donut represents the number of sequences acquired from that individual. **i,** Plot shows the number of germline nucleotide mismatches in the BCR sequences of clones isolated from the GC post-immunization. Data are compared using a Mann-Whitney test. **j,** IgGs were cloned from GC B cells 7 days after RBD immunization. ELISA traces show binding to RBD. C135 Fab high-affinity positive control (yellow) was included as a positive control.

**Supplementary data fig. 10:**
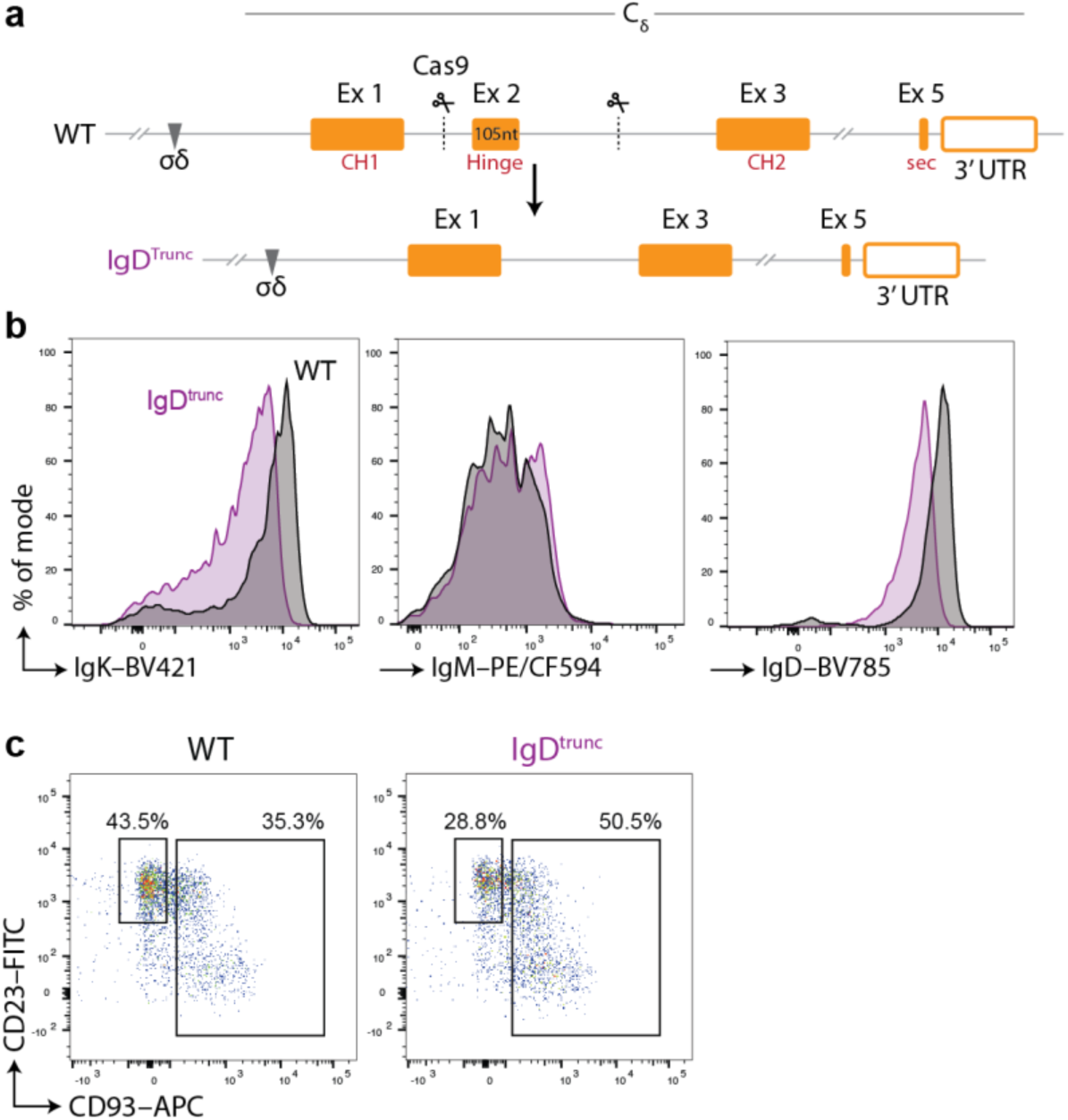
B cell development phenotyping of lgD^trunc^ mice. **a,** Graphical depiction of the C_δ_ region of the *Igh* loci and Cas9 gene segment excision strategy to produce the IgD^trunc^ allele by excising the exon 2, the symmetrical exon that encodes the discrete hinge portion of IgD. See methods section. **b,** Representative histograms showing the expression of IgM and IgD on B cells in the blood from both WT (black) and IgD^trunc^ mice. **c,** Representative flow cytometry plots showing the transitional and mature naïve B cell populations in mouse blood.

## References

1. Ohta, Y. & Flajnik, M. IgD, like IgM, is a primordial immunoglobulin class perpetuated in most jawed vertebrates. Proceedings of the National Academy of Sciences 103, 10723–10728 (2006).

2. Rowe, D. S. & Fahey, J. L. A New Class of Human Immunoglobulins. I. A Unique Myeloma Protein. J. Exp. Med. 121, 171–84 (1965).

3. Rowe, D. S., Crabbé, P. A. & Turner, M. W. Immunoglobulin D in serum, body fluids and lymphoid tissues. Clin. Exp. Immunol. 3, 477 (1968).

4. Yang, J. & Reth, M. Receptor Dissociation and B-Cell Activation. in 27–43 (2015). doi:10.1007/82_2015_482.

5. Venkitaraman, A. R., Williams, G. T., Dariavach, P. & Neuberger, M. S. The B-cell antigen receptor of the five immunoglobulin classes. Nature 352, 777–781 (1991).

6. Nemazee, D. Mechanisms of central tolerance for B cells. Nat. Rev. Immunol. 17, 281–294 (2017).

7. Cyster, J. G. & Allen, C. D. C. B Cell Responses: Cell Interaction Dynamics and Decisions. Cell 177, 524–540 (2019).

8. Hartley, S. B. et al. Elimination of self-reactive B lymphocytes proceeds in two stages: Arrested development and cell death. Cell 72, 325–335 (1993).

9. Hartley, S. B. et al. Elimination from peripheral lymphoid tissues of self-reactive B lymphocytes recognizing membrane-bound antigens. Nature 353, 765–769 (1991).

10. Nemazee, D. A. & Bürki, K. Clonal deletion of B lymphocytes in a transgenic mouse bearing anti-MHC class I antibody genes. Nature 337, 562–566 (1989).

11. Radic, M. Z., Erikson, J., Litwin, S. & Weigert, M. B lymphocytes may escape tolerance by revising their antigen receptors. J. Exp. Med. 177, 1165–1173 (1993).

12. Goodnow, C. C., Brink, R. & Adams, E. Breakdown of self-tolerance in anergic B lymphocytes. Nature 352, 532–536 (1991).

13. Tiegs, S. L., Russell, D. M. & Nemazee, D. Receptor editing in self-reactive bone marrow B cells. J. Exp. Med. 177, 1009–1020 (1993).

14. Victora, G. D. & Nussenzweig, M. C. Germinal Centers. Annu. Rev. Immunol. 40, 413–442 (2022).

15. Vitetta, E. S. et al. Cell surface immunoglobulin. XI. The appearance of an IgD-like molecule on murine lymphoid cells during ontogeny. J. Exp. Med. 141, 206–15 (1975).

16. Enders, A. et al. Zinc-finger protein ZFP318 is essential for expression of IgD, the alternatively spliced Igh product made by mature B lymphocytes. Proc. Natl. Acad. Sci. U. S. A. 111, 4513–4518 (2014).

17. Gutzeit, C., Chen, K. & Cerutti, A. The enigmatic function of IgD: some answers at last. Eur. J. Immunol. 48, 1101–1113 (2018).

18. Roes, J. & Rajewsky, K. Immunoglobulin D (IgD)-deficient mice reveal an auxiliary receptor function for IgD in antigen-mediated recruitment of B cells. J. Exp. Med. 177, 45–55 (1993).

19. Nitschke, L., Kosco, M. H., Köhler, G. & Lamers, M. C. Immunoglobulin D-deficient mice can mount normal immune responses to thymus-independent and -dependent antigens. Proc. Natl. Acad. Sci. U. S. A. 90, 1887–91 (1993).

20. Lutz, C. et al. IgD can largely substitute for loss of IgM function in B cells. Nature 393, 797–801 (1998).

21. Nechvatalova, J. et al. Absence of Surface IgD Does Not Impair Naive B Cell Homeostasis or Memory B Cell Formation in *IGHD* Haploinsufficient Humans. The Journal of Immunology 201, 1928–1935 (2018).

22. Noviski, M. et al. IgM and IgD B cell receptors differentially respond to endogenous antigens and control B cell fate. Elife 7, (2018).

23. Quách, T. D. et al. Anergic Responses Characterize a Large Fraction of Human Autoreactive Naive B Cells Expressing Low Levels of Surface IgM. The Journal of Immunology 186, 4640–4648 (2011).

24. Chen, Q., Menon, R., Calder, L. J., Tolar, P. & Rosenthal, P. B. Cryomicroscopy reveals the structural basis for a flexible hinge motion in the immunoglobulin M pentamer. Nat. Commun. 13, 6314 (2022).

25. Su, Q. et al. Cryo-EM structure of the human IgM B cell receptor. Science (1979). 377, 875–880 (2022).

26. Evans, R. et al. Protein complex prediction with AlphaFold-Multimer. Preprint at 10.1101/2021.10.04.463034 (2021).

27. Sun, Z. et al. Semi-extended Solution Structure of Human Myeloma Immunoglobulin D Determined by Constrained X-ray Scattering. J. Mol. Biol. 353, 155–173 (2005).

28. Davies, A. M. et al. Crystal structures of the human IgD Fab reveal insights into CH1 domain diversity. Mol. Immunol. 159, 28–37 (2023).

29. Depoil, D. et al. CD19 is essential for B cell activation by promoting B cell receptor–antigen microcluster formation in response to membrane-bound ligand. Nat. Immunol. 9, 63–72 (2008).

30. Treanor, B., Depoil, D., Bruckbauer, A. & Batista, F. D. Dynamic cortical actin remodeling by ERM proteins controls BCR microcluster organization and integrity. Journal of Experimental Medicine 208, 1055–1068 (2011).

31. Qi, S. Y., Groves, J. T. & Chakraborty, A. K. Synaptic pattern formation during cellular recognition. Proceedings of the National Academy of Sciences 98, 6548–6553 (2001).

32. Løset, G. A., Roux, K. H., Zhu, P., Michaelsen, T. E. & Sandlie, I. Differential Segmental Flexibility and Reach Dictate the Antigen Binding Mode of Chimeric IgD and IgM: Implications for the Function of the B Cell Receptor. The Journal of Immunology 172, 2925–2934 (2004).

33. Zhou, Z. et al. Genetically Encoded Short Peptide Tags for Orthogonal Protein Labeling by Sfp and AcpS Phosphopantetheinyl Transferases. ACS Chem. Biol. 2, 337–346 (2007).

34. Gregorio, G. G. et al. Single-molecule analysis of ligand efficacy in β2AR–G-protein activation. Nature 547, 68–73 (2017).

35. Steichen, J. M. et al. HIV Vaccine Design to Target Germline Precursors of Glycan-Dependent Broadly Neutralizing Antibodies. Immunity 45, 483–496 (2016).

36. Schaefer-Babajew, D. et al. Antibody-mediated diversification of primary and secondary humoral immune responses. Journal of Experimental Medicine 223, (2026).

37. Sabouri, Z. et al. IgD attenuates the IgM-induced anergy response in transitional and mature B cells. Nat. Commun. 7, 13381 (2016).

38. Übelhart, R. et al. Responsiveness of B cells is regulated by the hinge region of IgD. Nat. Immunol. 16, 534–543 (2015).

39. Chung, J. B., Silverman, M. & Monroe, J. G. Transitional B cells: step by step towards immune competence. Trends Immunol. 24, 342–348 (2003).

40. Goodnow, C. C., Adelstein, S. & Basten, A. The Need for Central and Peripheral Tolerance in the B Cell Repertoire. Science (1979). 248, 1373–1379 (1990).

41. Ota, T., Ota, M., Duong, B. H., Gavin, A. L. & Nemazee, D. Liver-expressed Igê superantigen induces tolerance of polyclonal B cells by clonal deletion not ê to λ receptor editing. Journal of Experimental Medicine 208, 617–629 (2011).

42. Wardemann, H. et al. Predominant Autoantibody Production by Early Human B Cell Precursors. Science (1979). 301, 1374–1377 (2003).

43. Russell, D. M. et al. Peripheral deletion of self-reactive B cells. Nature 354, 308–11 (1991).

44. Casellas, R. et al. Contribution of receptor editing to the antibody repertoire. Science 291, 1541–4 (2001).

45. Hägglöf, T. et al. Continuous germinal center invasion contributes to the diversity of the immune response. Cell 186, 147–161.e15 (2023).

46. Burnett, D. L., Reed, J. H., Christ, D. & Goodnow, C. C. Clonal redemption and clonal anergy as mechanisms to balance B cell tolerance and immunity. Immunol. Rev. 292, 61–75 (2019).

47. Dizon, B. L. P., Holla, P., Mutic, E. C., Schaughency, P. & Pierce, S. K. Human naïve B cells show evidence of anergy and clonal redemption following vaccination. NPJ Vaccines 10, 96 (2025).

48. Burnett, D. L. et al. Germinal center antibody mutation trajectories are determined by rapid self/foreign discrimination Europe PMC Funders Group. Science (1979). 360, 223–226 (2018).

49. Müller, R. et al. High-resolution structures of the IgM Fc domains reveal principles of its hexamer formation. Proc. Natl. Acad. Sci. U. S. A. 110, 10183–8 (2013).

50. Davis, A. C., Collins, C., Yoshimura, M. I., D’Agostaro, G. & Shulman, M. J. Mutations of the mouse mu H chain which prevent polymer assembly. J. Immunol. 143, 1352–7 (1989).

51. Booth, D. S., Avila-Sakar, A. & Cheng, Y. Visualizing Proteins and Macromolecular Complexes by Negative Stain EM: from Grid Preparation to Image Acquisition. Journal of Visualized Experiments 10.3791/3227 (2011) doi:10.3791/3227.

52. Mastronarde, D. N. Automated electron microscope tomography using robust prediction of specimen movements. J. Struct. Biol. 152, 36–51 (2005).

53. Juette, M. F. et al. Single-molecule imaging of non-equilibrium molecular ensembles on the millisecond timescale. Nat. Methods 13, 341–344 (2016).

54. Kiselev, R. et al. Parallel stopped-flow interrogation of diverse biological systems at the single-molecule scale. Nat. Methods 23, 78–87 (2026).

55. Senavirathne, G., Lopez, M. A., Messer, R., Fishel, R. & Yoder, K. E. Expression and purification of nuclease-free protocatechuate 3,4-dioxygenase for prolonged single-molecule fluorescence imaging. Anal. Biochem. 556, 78–84 (2018).

56. Robbiani, D. F. et al. Convergent antibody responses to SARS-CoV-2 in convalescent individuals. Nature 584, 437–442 (2020).

57. MacLean, A. J. et al. Affinity maturation of antibody responses is mediated by differential plasma cell proliferation. Science (1979). 387, 413–420 (2025).

58. Viant, C. et al. Antibody Affinity Shapes the Choice between Memory and Germinal Center B Cell Fates. Cell 183, 1298–1311.e11 (2020).

59. Gu, H., Zou, Y. R. & Rajewsky, K. Independent control of immunoglobulin switch recombination at individual switch regions evidenced through Cre-loxP-mediated gene targeting. Cell 73, 1155–64 (1993).

60. Russell, D. M. et al. Peripheral deletion of self-reactive B cells. Nature 354, 308–11 (1991).

61. Viant, C., Escolano, A., Chen, S. T. & Nussenzweig, M. C. Sequencing, cloning, and antigen binding analysis of monoclonal antibodies isolated from single mouse B cells. STAR Protoc. 2, 100389 (2021).

62. Martin, M. Cutadapt removes adapter sequences from high-throughput sequencing reads. EMBnet. J. 17, 10 (2011).

63. Dobin, A. et al. STAR: ultrafast universal RNA-seq aligner. Bioinformatics 29, 15–21 (2013).

64. Liao, Y., Smyth, G. K. & Shi, W. featureCounts: an efficient general purpose program for assigning sequence reads to genomic features. Bioinformatics 30, 923–930 (2014).

65. Hagemann-Jensen, M. et al. Single-cell RNA counting at allele and isoform resolution using Smart-seq3. Nat. Biotechnol. 38, 708–714 (2020).

66. Smith, T., Heger, A. & Sudbery, I. UMI-tools: modeling sequencing errors in Unique Molecular Identifiers to improve quantification accuracy. Genome Res. 27, 491–499 (2017).

67. Hao, Y. et al. Dictionary learning for integrative, multimodal and scalable single-cell analysis. Nat. Biotechnol. 42, 293–304 (2024).

68. Subramanian, A. et al. Gene set enrichment analysis: A knowledge-based approach for interpreting genome-wide expression profiles. Proceedings of the National Academy of Sciences 102, 15545–15550 (2005).

69. Liberzon, A. et al. Molecular signatures database (MSigDB) 3.0. Bioinformatics 27, 1739–1740 (2011).

70. Castanza, A. S. et al. Extending support for mouse data in the Molecular Signatures Database (MSigDB). Nat. Methods 20, 1619–1620 (2023).

71. Song, L. et al. TRUST4: immune repertoire reconstruction from bulk and single-cell RNA-seq data. Nat. Methods 18, 627–630 (2021).

72. Wang, Z. et al. Memory B cell development elicited by mRNA booster vaccinations in the elderly. J. Exp. Med. 220, (2023).

73. Deimel, L. P. et al. Clonal expansion and diversification of germinal center and memory B cell responses to booster immunization in primates. Cell Rep. 44, 116142 (2025).

74. Sonoda, E. et al. B Cell Development under the Condition of Allelic Inclusion. Immunity 6, 225–233 (1997).

